# Scaffold-mediated delivery of a miRNA-29b mimic mitigates excessive extracellular matrix deposition and matrix contraction in wound healing applications

**DOI:** 10.64898/2026.08.25.746629

**Authors:** Juan Carlos Palomeque Chávez, Amarachi Erugo, Marko Dobricic, Ahmed Al-Maini, Jack Maughan, James E. Dixon, Cathal J. Kearney, Shane Browne, Fergal J. O’Brien

**Affiliations:** Tissue Engineering Research Group, Department of Anatomy & Regenerative Medicine, Royal College of Surgeons in Ireland, Dublin, Ireland; Advanced Materials and Bioengineering Research Centre (AMBER), Royal College of Surgeons in Ireland and Trinity College Dublin, Dublin, Ireland; Kearney Lab, Department of Biomedical Engineering, University of Massachusetts, Amherst, USA; Regenerative Medicine & Cellular Therapies (RMCT), Biodiscovery Institute (BDI), School of Pharmacy, University of Nottingham, Nottingham, United Kingdom; NIHR Nottingham Biomedical Research Centre, University of Nottingham, Nottingham, United Kingdom; Trinity Centre for Biomedical Engineering, Trinity College Dublin, Dublin, Ireland; Centre for Research in Medical Devices (CÚRAM), University of Galway, Galway, Ireland

**Keywords:** gene delivery, miRNA, collagen, fibrosis, wound healing

## Abstract

Disruption of the wound healing cascade can result in pathological outcomes, including fibrosis due to myofibroblast-mediated contraction and collagen deposition. Despite the clinical significance, effective treatments for fibrosis remain limited as current therapies often show inconsistent efficacy, adverse effects, and patient discomfort. Combinatorial therapeutic strategies integrating biomaterial scaffolds with gene delivery have shown promise in regenerative healing. MicroRNAs (miRNAs) are key regulators of fibrotic signalling in cells, including fibroblasts and myofibroblasts. Specifically, miRNA-29b is notable for downregulating pro-fibrotic genes, including collagen type I, reducing ECM accumulation, and limiting fibroblast/myofibroblast overactivation. In this context, the present work develops a collagen-GAG (CG) scaffold platform for delivery of miRNA-29b complexed GET nanoparticles to inhibit fibrosis. Initially, bioinformatic analysis of miRNA-29b validated its involvement in ECM-associated pathways and processes, followed by successful nanoparticle internationalisation in primary dermal fibroblasts. The anti-fibrotic efficacy of the optimised miRNA-29b nanoparticles was subsequently demonstrated by significant reductions in collagen deposition and α-SMA expression, both key indicators of myofibroblast differentiation and fibrosis. The optimised miRNA-29b formulation was then incorporated into 3D porous collagen-GAG (CG) scaffolds, which modulated fibrotic gene expression while preserving scaffold structure conducive to fibroblast/myofibroblast infiltration and proliferation. Finally, functional outcomes of seeded TGF-β-stimulated fibroblasts, including reduced matrix contraction, α-SMA expression, and ECM deposition, were comparable to those observed in non-fibrotic conditions, thereby confirming the therapeutic potential of scaffold-mediated miRNA- 29b delivery. Together, these findings demonstrate that scaffold-mediated miRNA-29b delivery represents a promising anti-fibrotic strategy for wound healing by mitigating myofibroblast activation, limiting matrix contraction, and preventing pathological ECM accumulation.

**Highlights:**

- miRNA-29b delivery mitigates fibrosis via collagen and α-SMA attenuation
- Scaffold-mediated miRNA-29b delivery inhibits fibrosis-related matrix contraction
- Scaffold-mediated miRNA-29b delivery mitigates excessive extracellular matrix deposition

**Graphical Abstract:** 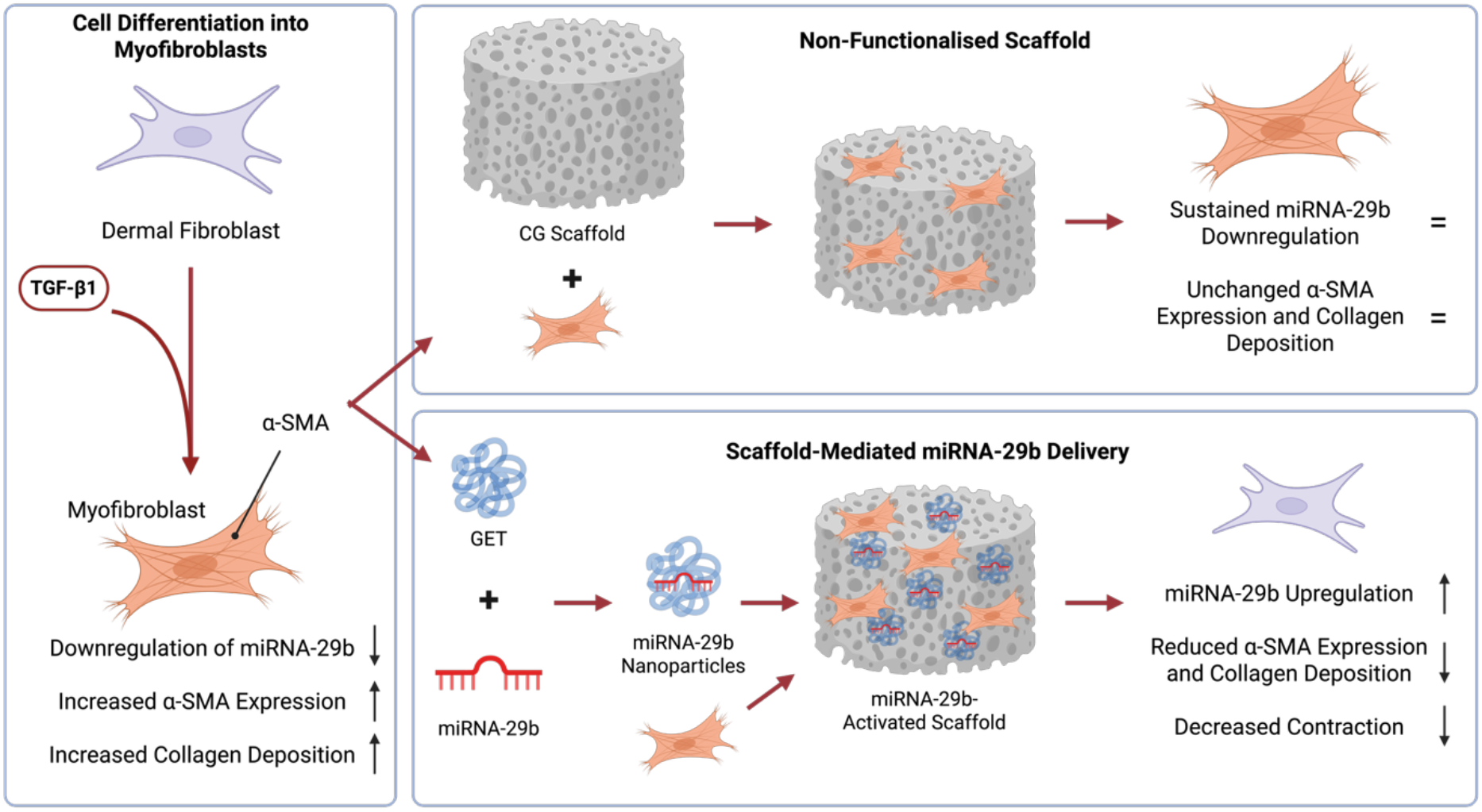

## 1. Introduction

Wound healing is initiated following skin injury and, under healthy normal physiological conditions, culminates in wound closure and restoration of the skin’s homeostatic functions. Disruption at any stage of this process can result in pathological outcomes, including fibrosis.^1–3^ Fibrotic wound healing is characterised by persistent myofibroblast activation, excessive ECM accumulation, and aberrant scarring, leading to both functional impairments and aesthetic burden.^4,5^ Clinically, these pathological processes are exemplified by hypertrophic scars and keloids, which vary in incidence and prevalence across different populations.^6,7^ Despite their significant clinical and socioeconomic impact, effective treatment options remain limited. Current interventions, including silicone sheeting, corticosteroid injections, and cryotherapy, are widely used but often provide inconsistent clinical benefit and are associated with adverse side effects and patient discomfort.^8–10^ These limitations underscore the urgent need for more effective, targeted anti-fibrotic therapies that address the underlying molecular mechanisms driving pathological wound repair.

At the molecular level, fibrosis during wound healing is primarily driven by the transforming growth factor-beta (TGF-β) signalling pathway, a central regulator of fibroblast activation and myofibroblast differentiation.^11^ TGF-β promotes the contractile myofibroblast phenotype, marked by increased expression of alpha-smooth muscle actin (α-SMA). Canonical TGF-β signalling is mediated through SMAD2 and SMAD3 phosphorylation, which subsequently form a complex with SMAD4 and translocate to the nucleus to regulate the transcription of pro-fibrotic genes, such as collagen type I (*COL1A1*) and α-SMA (*ACTA2*).This pathway is tightly controlled by the inhibitory protein SMAD7, which prevents SMAD2/3 phosphorylation and thereby suppresses downstream transcriptional activity^12,13^ However, pro-inflammatory cytokines can modulate canonical TGF-β/SMAD signalling through non-canonical mechanisms, including downregulation of SMAD7 and activation of pathways such as STAT3, NF- κB, and MAPKs.^14–18^ In chronic or dysregulated wounds, pro-inflammatory (M1) macrophages further exacerbate fibrosis through sustained secretion of inflammatory cytokines and proteolytic enzymes, prolonging inflammation and perpetuating myofibroblast activation.^19,20^ These interconnected mechanisms underscore the need for anti-fibrotic therapies capable of targeting fibrosis across diverse wound aetiologies.

Achieving an appropriate balance in fibroblast activation is therefore essential for effective wound healing. Insufficient activation delays ECM deposition and wound closure, whereas overactivation leads to pathological scarring and compromised tissue function.^4,5^ To address this challenge, combinatorial therapeutic strategies that integrate biomaterial scaffolds with gene delivery have emerged as promising tools to enhance regenerative healing.^21–23^ Among these, collagen-based biomaterials are widely used in the treatment of wounds of diverse aetiologies, with several collagen-based wound dressings already established in clinical practice.^24–27^ These scaffolds provide a structural template that facilitates cell infiltration, supports tissue remodelling, and enhances the overall wound healing response. Importantly, their ease of functionalisation enables the incorporation of advanced therapeutic strategies, including localised gene delivery, to further promote tissue regeneration.^28^ Our group has extensive experience developing collagen-glycosaminoglycan (CG) scaffolds for regenerative wound healing applications.^21,28,29^ These scaffolds have been shown to reduce wound contraction while promoting tissue regeneration and providing an effective platform for the local delivery of nucleic acid-based therapeutics.^28,30^ By enabling spatially controlled modulation of gene expression within the wound microenvironment, gene-activated scaffolds offer a promising strategy to regulate fibroblast behaviour, attenuate fibrosis, and minimise excessive matrix contraction while preserving normal tissue repair.

RNA interference (RNAi) strategies have attracted considerable interest for the functionalisation of biomaterial scaffolds. RNAi is a naturally occurring cellular mechanism that operates in the cytoplasm, where gene silencing is achieved through the degradation of messenger RNA (mRNA) or the repression of its translation.^31,32^ Among RNAi molecules, microRNAs (miRNAs) are small non-coding RNAs that regulate cell behaviour via the post-transcriptional modulation of multiple target genes.^3,33^ Several miRNAs have been identified as key regulators of fibroblast activation, ECM remodelling, and the TGF-β/SMAD signalling, underscoring their potential as anti-fibrotic therapeutics.^34^ Among these, miRNA-29b has emerged as a particularly promising candidate owing to its ability to suppress the expression of pro- fibrotic genes such as collagen type I, thereby reducing ECM accumulation and limiting myofibroblast activation.^35–37^ Beyond cutaneous wound healing, miRNA-29b has also been implicated in the regulation of renal^38^, pulmonary^39^, and hepatic^40^ fibrosis, further highlighting its broad therapeutic relevance.

The clinical translation of nucleic acid-based therapeutics remains challenging due to poor cellular uptake and rapid degradation of free nucleic acids under physiological conditions, highlighting the appropriate selection of vector to ensure cargo delivery.^41,42^ While viral vectors offer high transfection efficiency, non-viral delivery systems present advantages in terms of scalability and reduced immunogenicity.^31,42^ In this context, the non-viral glycosaminoglycan enhanced transduction (GET) peptide has demonstrated enhanced cellular uptake across multiple cell types by combining cell penetrating and heparan sulphate GAG-binding domains, making it a promising vector for miRNA-29b delivery. ^43–45^

Building on these principles, the present study aimed to develop a CG scaffold-based platform for delivery of miRNA-29b complexed GET nanoparticles to reduce myofibroblast activation, prevent matrix contraction, and inhibit excessive ECM accumulation in wound healing. First, the role of miRNA-29b in regulating ECM-associated pathways was validated through bioinformatic analyses, followed by confirmation of efficient fibroblast internationalisation. The anti-fibrotic activity of the optimised miRNA-29b nanoparticles was subsequently demonstrated by significant reductions in collagen deposition and α-SMA expression in TGF-β1-stimulated fibroblasts, two established markers of fibrosis. Following incorporation into collagen-GAG (CG) scaffolds, the optimised miRNA-29b formulation effectively modulated fibrotic gene expression while preserving the porous scaffold architecture conducive to fibroblast infiltration and proliferation. Importantly, when seeded with TGF-β1- stimulated fibroblasts, scaffold-mediated delivery of miRNA-29b significantly reduced matrix contraction, α-SMA expression, and ECM deposition, restoring these functional outcomes to levels comparable with those observed under non-fibrotic conditions. Collectively, these findings demonstrate that scaffold-mediated miRNA-29b delivery represents a promising anti-fibrotic strategy for wound healing by mitigating myofibroblast activation, limiting matrix contraction, and preventing pathological ECM accumulation.

## 2. Materials and Methods

All reagents were purchased from Thermo Fisher Scientific (Ireland) unless otherwise stated. All cell culture was performed at 37°C and 5% CO_2_ unless otherwise stated.

### 2.1. Predictive pathway analysis of miRNA-29b interactions using publicly available datasets

To identify pathway interactions of miRNA-29b and evaluate its impact on ECM-associated mechanisms, we performed in silico analysis of miRNA interactions using publicly available datasets (TargetScan, miRDB, and StarBase). Predicted targets of human miRNA-29b-3p were retrieved from these databases, and the resulting target lists were intersected to generate a consolidated overall set of miRNA-29b targets (116 genes). The gene set was subsequently analysed using the online Database for Annotation, Visualization and Integrated Discovery (DAVID). Enriched Gene Ontology (GO) biological processes and Kyoto Encyclopaedia of Genes and Genomes (KEGG) pathways associated with ECM regulation and interactions were then examined to support the selection of miRNA-29b as a potential anti-fibrotic therapeutic target.

### 2.2. Assessment of ɑ-SMA expression in TGF-β1-stimulated fibroblasts

Normal human dermal fibroblasts (HDFs) were purchased from PromoCell (Germany) and culture in growth media containing low glucose (1.0 g/L) Dulbecco’s Modified Eagles Medium (DMEM) supplemented with 10% FBS and 1% penicillin-streptomycin (P/S). To induce myofibroblast differentiation, HDFs were seeded on tissue culture plates and cultured with growth medium containing transforming growth factor-beta 1 (TGF-β1, Cat# 100-21) at various concentrations for 24h. After this, media was removed and replaced with normal growth medium and collected at different timepoints.

Cell morphology, distribution, and α-SMA expression in myofibroblast/fibroblast populations were analysed through immunofluorescence staining. Initially, TGF-β1- conditioned HDFs were seeded on 13 mm round coverslips (Cat# 17274914, Fisher Scientific, UK). Following culture, cells were fixed in 4% PFA for 1h at 4°C before being washed 3 times with DPBS and stored at 4°C until use. Cells were permeabilised with 0.1% Triton X-100 solution for 5 min followed by incubation with 1% bovine serum albumin (BSA) for 2h to block the samples.

Following blocking, cells were incubated with anti-ɑ-SMA antibody (1:200, Cat# 14- 9760-82) overnight at 4°C. Then, cells were incubated with a DPBS solution containing Alexa Fluor 555 Phalloidin™ (1:500) and anti-mouse secondary antibody Alexa Fluor™ 488 (1:1000, Cat# A-21050) for 2 hours and Hoechst 33342 (1:10000) for 15 min with three DPBS washes in between steps. Finally, cells on coverslips were mounted on glass slides with Fluoromount-G™ Mounting Medium (Cat# 00-1958-02). All coverslips were imaged with a Nikon Eclipse 90i fluorescent microscope, maintaining gain, exposure, and magnification constant. Images were analysed using FIJI software^46^ to calculate nuclei count, cell coverage, and ɑ-SMA expression using automated scripts

### 2.3. Development, characterisation, and uptake analysis of miRNA nanoparticles

#### Formulation and physicochemical characterisation of miRNA nanoparticles

The miRIDIAN microRNA mimic hsa-miRNA-29b-3p (miRNA-29b) and scramble mimic negative control (miRNA-Scr) (Dharmacon, UK) were combined with the positively charged glycosaminoglycan enhanced transduction (GET) peptide through electrostatic interactions to form complexes at charge ratio 1:4 (CR4), 1:6 (CR6), and 1:8 (CR8).^21^ Additionally, miRIDIAN microRNA mimic red transfection control (Dharmacon, UK) was complexed with GET at CR8 to enable fluorescent tracking of nanoparticles (Cy3 NPs).

Physicochemical characterisation of the miRNA-29b and miRNA-Scr nanoparticles was carried out by dynamic light scattering (DLS) (Zetasizer 3000 HS, Malvern, UK) and nanoparticle tracking analysis (NTA) (NanoSight NS300, Malvern, UK) to determine charge (zeta potential) and size distribution as previously reported.^21^ Briefly, miRNA-29b nanoparticles were prepared with molecular grade water (MG-H_2_O). The volume was then increased to 1 mL and transferred to a disposable folded capillary cell (Malvern, UK) before DLS analysis. For NTA assessment, data was captured with sCMOS camera and a Blue488 laser, with data evaluation being carried out with the NTA 2.3 software (Malvern, UK).

#### Uptake analysis of miRNA nanoparticles

To assess the internalisation dynamics of miRNA-29b nanoparticles within HDFs, 4.8 x 10^3^ cells were seeded on a 12 well μ-slide (Cat#81201, ibidi, Germany) at a concentration of 1.92 x 10^4^ cells mL^-1^. Simultaneously, Cy3 NPs (20 pmol) were added to the cells to track nanoparticle uptake and distribution. The slide was then transferred to a Zeiss Celldiscoverer 7 microscope and images were taken every 30 min for 96h. After culture and imaging, cells were fixed and permeabilised as previously stated before staining with Alexa Fluor 488 Phalloidin™ (1:1000, Cat# A12379) for 1h and Hoechst 33342 (1:10000) for 15 min. Images were then taken with a Nikon Eclipse 90i fluorescent microscope. All image analysis and processing were then carried out with Fiji software.

### 2.4. Assessment of cell viability and gene expression post-transfection

#### Assessment of cell viability post-transfection

Following dermal fibroblast transfection with miRNA-29b nanoparticles, cell metabolic activity and DNA content were determined through Alamar Blue™ Cell Viability assay and Quant-iT™ PicoGreen™ dsDNA Assay according to the manufacturer’s protocol, respectively. Briefly, media was removed and growth medium containing 10% Alamar Blue™ reagent was added to the cells (500 μL). Samples were then incubated for 1h at 37°C. The supernatant was then collected, and the fluorescence of each sample was measured in triplicate (ex: 570 nm, em: 585 nm) using an Infinite 200 PRO plate reader (Tecan Group Ltd., Switzerland). Fluorescence measurements of the transfected groups were normalised in relation to the un-transfected control.

DNA content was determined through Quant-iT™ PicoGreen™ dsDNA Assay according to the manufacturer’s protocol. Media was removed and wells were flooded with 1 mL buffer (0.2 M sodium carbonate + 0.1% Triton X-100 in DI H_2_O) to lyse the cells. Samples were then subjected to 3 freeze-thawing cycles at −80°C before measurements were carried out. Fluorescence measurements (ex: 480 nm, em: 520 nm) were performed using an Infinite® 200 PRO plate reader. Finally, the DNA concentration was extrapolated from the standard curve.

#### Assessment of gene expression post-transfection via qRT-PCR

RNA expression was assessed through quantitative real time PCR (qRT-PCR). On day 3 post-transfection, RNA was extracted from fibroblasts by lysing cells using 500 μL QIAzol lysis reagent (Qiagen, Ireland) per sample and subsequently extracting the RNA fraction using a miRNeasy kit (Qiagen, Ireland) according to the manufacturer’s protocol. cDNA templates were then produced for RNA and microRNA analysis using a QuantiTect Reverse Trancription Kit (Qiagen, Ireland) and a Taqman™ Advanced miRNA cDNA synthesis kit, respectively. qRT-PCR was carried out with a variety of target genes (Table 4-1) using a Lightcycler 480 II (Roche, UK). Finally, the ΔΔCt method was utilised to calculate the fold change expression of the genes of interest.

### 2.5. Intracellular protein expression analysis following miRNA-29b transfection

To determine *phospho-*SMAD2/3, α-SMA, and vinculin protein expression levels following miRNA-29b transfection in HDFs, western blots were conducted. Proteins were extracted using RIPA buffer (Cat# 89901), followed by measurement of the total protein content using a Pierce™ BCA Protein Assay Kit (Cat# 23225). Cell lysates were then centrifuged at 10000 g for 10 min. Following this, 9 μg total protein was incubated at 95°C with loading buffer (Cat# 161-0747, BioRad) before loading into gradient gels (Mini-PROTEAN TGX Gels 4-20%, BioRad). Proteins were separated by electrophoresis and subsequently transferred to polyvinylidene fluoride (Trans-Blot Turbo Transfer – 0.2 µm PVDF, BioRad) membranes and blocked with 5% (w/v) fat-free milk in 0.1% (v/v) Tris-buffered saline – Tween 20 (TBST). Then, membranes were incubated overnight at 4°C in a 5% BSA solution in TBST containing 1:1000 dilution of anti-*phospho-*SMAD2/3 (Cat# 8828, Cell Signalling Technologies), anti-α-SMA (Cat# 19245, Cell Signalling Technologies), or anti-vinculin (Cat#13901, Cell Signalling Technologies) antibodies. Membranes were washed three times in TBST before incubation with secondary antibody (1:10000) for 1 h at RT. Protein bands were visualised using an Amersham Imager 600.

Following visualisation and image collection, membranes were stripped of antibodies using the Restore™ Plus Western Blot Stripping Buffer (Cat# 46430) according to the manufacturers protocol. Then, membranes were washed and blocked as previously stated before overnight incubation with anti-GAPDH (Cat# 5174, Cell Signalling Technologies) antibody at 4°C. Similarly, membranes were incubated with the secondary antibody and imaged using an Amersham Imager 600. Relative protein expression was normalised to GAPDH expression for every group.

### 2.6. Assessment of functional outcomes post-transfection of miRNA-29b

#### Quantification of collagen deposition via histological staining

To assess the effect of miRNA-29b delivery to HDFs on collagen deposition, cells were plated on coverslips, conditioned with TGF-β1, and transfected as before. Additionally, L-ascorbic acid (50 µg mL^-1^, Cat# A7506, Merck) was supplemented in the growth medium to stabilise collagen deposition. Following cell culture, cells were fixed with 4% PFA for 15 min at RT before being washed 3 times with DPBS and stored at 4°C. Then, samples were dehydrated, stained with picrosirius red to visualise collagen, and rehydrated. Finally, stained coverslips were mounted overnight on glass slides with DPX mountant for histology (Cat# 06522, Merck, Ireland) before simultaneous bright- field and fluorescent imaging with a Nikon Eclipse 90i microscope, maintaining consistent magnification, gain, and exposure. Final processing and quantification was carried out in Fiji software.

#### Assessment of adhesion protein co-localisation through immunofluorescence

Co-localisation and relative expression of vinculin and α-SMA in myofibroblast/fibroblast populations were analysed through immunofluorescence staining. Initially, TGF-β1-conditioned HDFs were seeded on coverslips. Following culture, cells were fixed, permeabilised, and blocked as previously stated. Then, cells were incubated with anti-α-SMA (1:200, Cat# 701457) and anti-vinculin eFluor™ 570 (1:200, Cat# 41-9777-82) antibodies overnight at 4°C. Samples were incubated with anti-rabbit secondary antibody Alexa Fluor™ 488 (1:1000, Cat# A-11008) for 1 h and Hoechst 33342 (1:10000) for 15 min with three DPBS washes between steps. Finally, cells on coverslips were mounted on glass slides with Fluoromount-G™ Mounting Medium. All coverslips were imaged using a Zeiss LSM 710 confocal microscope, maintaining consistent gain, exposure, and magnification. Images were analysed using FIJI software to calculate nuclei count and relative expression of α-SMA and vinculin using automated scripts.

### 2.7. Collagen-GAG (CG) scaffold fabrication, miRNA functionalisation, and release profile

The CG slurry used for scaffold fabrication was prepared as previously described.^47^ Briefly, a solution combining 0.5% w/v of microfibrillar type I collagen (Integra Life Sciences, USA) and 0.05% w/v chondroitin-6-sulfate (GAG) (Sigma-Aldrich, Germany) was prepared using acetic acid (0.05M) as a solvent. The solution was blended using an Ultra-Turrax® T25 homogenizer (IKA, Germany) at 15,000 rpm at 4°C. Then, the slurry was degassed under vacuum (∼5 Torr) at room temperature before being stored at 4°C until use. CG scaffolds were prepared by lyophilization process by pipetting 400 µL of into 10 mm diameter stainless steel moulds before freezing to a final temperature of −10°C, reduced at a rate of 1°C min^-1^ and maintained for 60 min. The solvent was sublimated under vacuum (200 mTorr) for 24h at 0°C.

To enhance their structural properties, CG scaffolds were chemically crosslinked with 1-ethyl-3-(3-dimethyl aminopropyl)-carbodiimide (EDAC) and N-hydroxy succinimide (NHS) (Sigma-Aldrich, Germany) as previously described.^47^ Briefly, scaffolds were hydrated in DPBS for 1h prior to crosslinking. DI H_2_O solutions containing EDAC (6 mmol per gram of collagen) and NHS (2.5 M ratio of EDAC:NHS) were prepared and mixed. Scaffolds were then transferred to the EDAC/NHS solutions and allowed to crosslink for 2h at room temperature. After crosslinking, scaffolds were washed before sterilization in 70% ethanol and washed under sterile conditions.

CG scaffolds were then gene-activated through the soak-loading of miRNA-29b mimic (CG-29b) or miRNA-Scr mimic (CG-Scr) nanoparticles. Briefly, miRNA nanoparticles were prepared as previously described and soak-loaded on one side of the CG scaffolds (40 pmol miRNA) and incubated for 45 min at 37°C. Then, scaffolds were flipped, and the process was repeated. After soak-loading the miRNA nanoparticles, 1.25 x 10^5^ hDFs were seeded onto the first side of the scaffolds at a concentration of 2.5 x 10^6^ cells mL^-1^. Scaffolds were allowed to incubate at 37°C for 15 mins before repeating the process on the opposite side. Finally, growth medium containing transforming growth factor-beta 1 (10 ng mL^-1^, TGF-β1, Cat# 100-21) was added to the wells for 24h. Then, media was removed and replaced with normal growth medium and collected at different timepoints.

To confirm that functionalisation of CG scaffolds with miRNA nanoparticles did not affect the microstructure, scaffolds were imaged using a scanning electron microscope (SEM) as previously described.^48^ Briefly, scaffolds were gene-activated before being submerged in ethanol in preparation for supercritical CO_2_ drying which was performed using a K850 Critical Point Dryer (Quorum Tech, UK) with a bleed time of 1h. Dried scaffolds were bisected using a scalpel, followed by mounting on metallic pin studs with carbon tape. Mounted scaffolds were sputtered with an 80/20 mixture of gold/palladium alloy to a thickness of ∼4 nm in a Cressington 108 auto sputter coater. Scaffold microstructure was assessed using a Zeiss Ultra FE-SEM (Zeiss, Germany) with an accelerating voltage of 3kV at several magnifications.

To assess the miRNA release profiles from miRNA-activated scaffolds, miRNA-29b nanoparticles were formulated in MG H_2_O before soak loading on scaffolds (0.2 nmol). Samples were then placed in 24-well plates and flooded with 2 mL MG H_2_O. The release assessment was carried out under static conditions at 37°C. At every timepoint, 200 μL of supernatant were collected and replaced with 200 μL of fresh MG H_2_O. The supernatant containing released nanoparticles was then incubated with 50 μL of heparin (1 mg mL^-1^) for 90 min to promote nanoparticle dissociation. Finally, the amount of miRNA released was quantified using a Quant-it™ RiboGreen Reagent and RNA Assay kit following the manufacturer’s protocol.

### 2.8. Assessment of anti-fibrotic responses from TGF-β1-stimulated dermal fibroblasts on CG-29b scaffolds

To understand the effect of scaffold-mediated miRNA-29b transfection on the viability of HDFs, metabolic activity was analysed through an Alamar Blue™ Cell Viability assay. The assay was carried as previously described with 10% Alamar Blue-containing growth medium (1 mL) and an incubation for 2h. Fluorescence measurements of the miRNA transfected groups were normalised to the gene-free (CG) control.

To assess the anti-fibrotic response following scaffold-mediated miRNA-29b delivery to HDFs through the modulation of α-SMA expression, scaffolds were stained with immunofluorescent probes. Following cell culture, scaffolds were fixed in 4% PFA for 1h at 4°C before being permeabilised and blocked as previously described. Scaffolds were then incubated with anti-ɑ-SMA antibody overnight at 4°C, followed by staining with Alexa Fluor 555 Phalloidin™ (1:500), secondary antibody Alexa Fluor™ 488 (1:1000), and Hoechst 33342 (1:10000). Finally, scaffolds were imaged using a Zeiss LSM 710 confocal microscope, maintaining consistent gain, exposure, and magnification. Images were analysed using FIJI software to calculate nuclei count, cell coverage, and α-SMA expression using automated scripts. RNA expression was assessed through qRT-PCR on day 3 post-transfection as previously described.

To assess the level of contraction inhibition in myofibroblast/fibroblast cell populations following miRNA-29b treatments, a functional gel contraction assay was carried out. miRNA nanoparticles were prepared as previously described (0.12 nmol miRNA, 50 μL total volume) and mixed with 150 μL of HDFs suspension at a concentration of 5.0 x 10^5^ cells mL^-1^. Finally, 100 μL of collagen I rat tail solution (Cat# A1048301) were mixed with the cell/nanoparticles suspension to create a 1 mg mL^-1^ collagen gel. Solutions were pipetted into 10 mm round steel moulds and incubated at 37°C for 30 min to allow gels to polymerise. Following incubation, gel discs were released from moulds and transferred to 6-well plates and cultured in TGF-β1-containing growth medium for 24h before transferring to normal growth medium. Images of gels were taken at 1, 3, 5 and 7 days using a Leica EZ4W stereo microscope, maintaining consistent magnification. Image analysis and processing was carried out in Fiji.

To characterise the cell-associated ECM deposition dynamics following scaffold- mediated miRNA-29b delivery, the metabolic labelling method previously reported by Loebel et al^49^ was used. Briefly, dermal fibroblasts were cultured and expanded in standard growth medium until before seeding on CG scaffolds as mentioned previously. Following trypsinisation, cells were resuspended and cultured in glutamine-, methionine-, and cysteine-free high glucose DMEM supplemented with 10% FBS, 1% P/S, 50 μg mL^-1^ L-ascorbic acid, 0.201 mM L-cysteine, 0.1 mM L- azidohomoalanine (AHA), and 100 μM L-methionine. Medium was also supplemented with 10 ng mL^-1^ TGF-β1 (for supplemented groups) for 24 h before returning to TGF- β1-free medium afterwards.

After culture, scaffolds were fixed and deposited ECM labelling was performed by incubating the samples with 2% BSA solution containing 30 μM DBCO-488 (Cat # CCT-1278, 2BScientific) for 20 min at 37°C. Scaffold were then washed with PBS before being permeabilised with 0.1% Triton X-100 for 5 min and blocked with 1% BSA for 2h. Samples were incubated with anti-vimentin antibody (1:200, Cat #MA5-16409) overnight at 4°C before adding a PBS solution containing Alexa Fluor 555 Phalloidin™ (1:500) and secondary antibody anti-rabbit Alexa Fluor™ 647 (1:1000, Cat # A-21245) for 2h at RT. Nuclei were counterstained with Hoechst 33342 (1:10000) for 15 min. Finally, scaffolds were imaged using a Zeiss LSM 900 Airyscan confocal microscope, maintaining consistent gain, exposure, and magnification. Images were analysed using FIJI software to calculate vimentin expression, ECM coverage and ECM intensity using automated scripts.

### 2.9. Statistical Analysis

Statistical analysis was carried out using Graph-Pad Prism v 11.0.2. One-way ANOVA with a Tukey post-hoc test was used when more than one treatment was compared. Two-way ANOVA with Bonferroni post-hoc test was used when more than one treatment was compared across two factors. Results are expressed as mean ± standard deviation (SD), * indicates p<0.05, ** p<0.01, *** p<0.001, and *** p<0.001 in all instances.

## 3. Results

### 3.1. *In vitro* and *in silico* analyses confirm the efficient internalisation of miRNA-29b and its involvement in ECM-associated processes

Non-coding microRNAs (miRNAs) have the ability to post-transcriptionally modulate a cassette of genes via RNA interference mechanisms leading to multiple cellular responses.^3,33^ However, further characterisation is required to ensure the target group of genes is targeted to influence the desired biological outcome. To ensure the involvement of miRNA-29b in the modulation of ECM-related pathways, predictive bioinformatic analysis was carried out on target genes of miRNA-29b highlighting 116 common genes across 3 publicly available databases (Fig. 1A). Gene ontology assessment of predicted miRNA-29b target genes underscored the enrichment of processes associated with ECM organisation and assembly (Fig. 1B). Similarly, KEGG pathway analysis highlighted the enrichment of canonical and non-canonical pathways involved in ECM deposition and remodelling in fibroblasts, such as WNT^50^ and PI3K- AKT^51^ signalling pathways (Fig. 1C).

**Figure 1.**
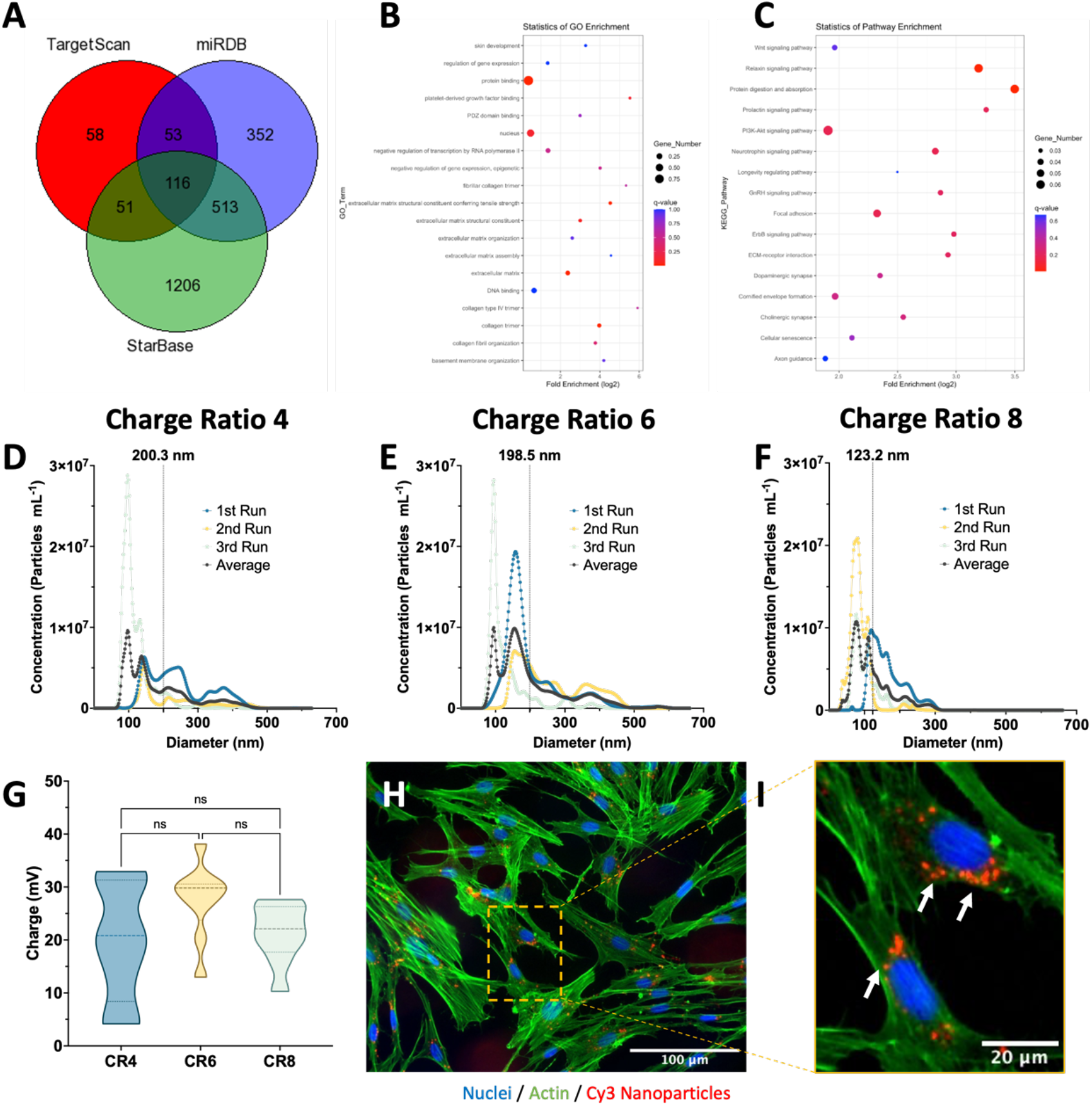
Predictive bioinformatic analysis and physicochemical characterisation of miRNA-29b provides evidence of optimal properties for fibroblast internalisation. (n=3) A) Venn diagram of predicted miRNA-29b targets identified across three public miRNA–mRNA interaction databases, showing the gene set used for enrichment analysis. B-C) Gene Ontology (GO) and KEGG enrichment analysis of common predicted miR-29b targets, highlighting the involvement of ECM-associated processes and pathways. D-G) Characterisation of miR-29b nanoparticles’ size distribution at CR4, CR6 and CR8 indicates a decreasing trend in diameter with increasing CR without affecting overall charge. H-I) Fluorescent imaging reveals successful nanoparticle internalisation as observed by perinuclear co-localisation of Cy-3-tagged nanoparticles within cells. ns indicates a non-significant difference. (n=3)

Having confirmed the involvement of miRNA-29b in ECM-associated processes, we assessed the physicochemical properties of miRNA-29b nanoparticles and their successful internalisation within human dermal fibroblasts. GET-complexed miRNA- 29b mimic nanoparticles at charge ratios (CR) 4, 6, and 8 exhibited a decreasing trend in average diameter with increasing GET peptide content. CR4 nanoparticles showed the biggest diameter at 200.3 ± 99.3 nm (Fig. 1D), followed by CR6 nanoparticles at 198.5 ± 85.7 nm (Fig. 1E), and CR8 nanoparticles at 123.2 ± 56.5 (Fig. 1F). Subsequent analysis of charge through dynamic light scattering (DLS) revealed that, despite the differences in size, all nanoparticle formulations presented an average positive charge of 20 mV or higher (Fig. 1G), necessary for successful cellular uptake. Additionally, fluorescent imaging of tagged nanoparticles in culture with dermal fibroblasts (Fig. 1H) confirmed nanoparticle uptake as highlighted by perinuclear colocalization of the nanoparticles within the cell cytoplasm (Fig. 1I).

### 3.2. Efficient anti-fibrotic responses from myofibroblasts following miRNA- 29b delivery are charge ratio and dose-dependent without compromising cell function

Following characterisation of physicochemical properties, nanoparticle uptake was evaluated by live-cell imaging of Cy3-labelled miRNA nanoparticles in dermal fibroblasts. Importantly, treatment with the miRNA-loaded nanoparticles preserved the characteristic spindle-shaped morphology of dermal fibroblasts (Fig. 2A), comparable to untreated controls (Supp. Fig. 1). Cells maintained normal migratory and proliferative behaviour despite continuous nanoparticle uptake for up to 4 days (Fig 2B), indicating that the delivery system did not induce cytotoxicity or impaired cellular function. Notably, the Cy3-tagged miRNA remained detectable in both daughter cells following cell division, demonstrating stable intracellular retention while further supporting the safety and biocompatibility of the nanoparticle delivery platform.

**Figure 2.**
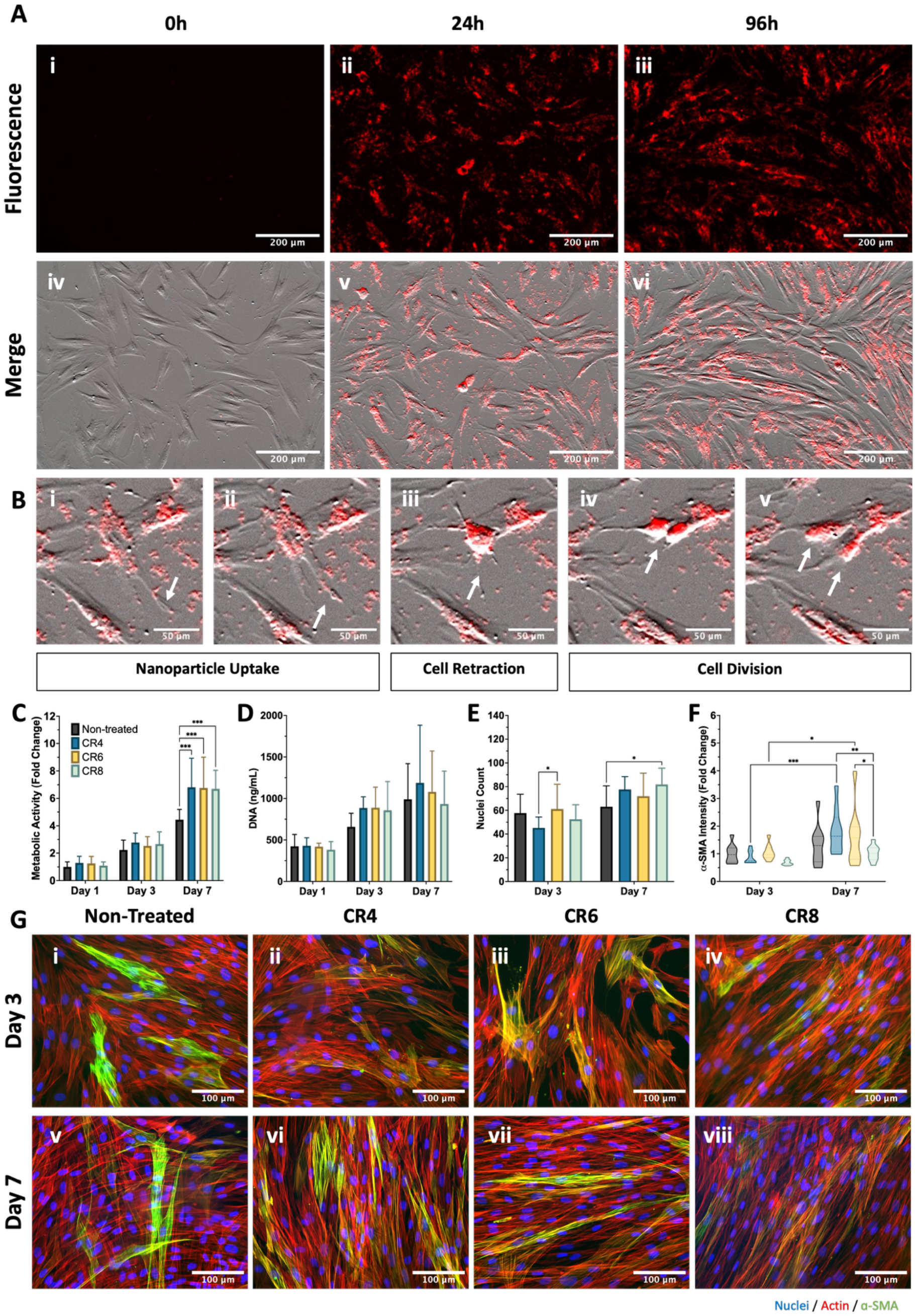
Efficient intracellular delivery of optimised miRNA-29b nanoparticles demonstrates strong anti-fibrotic activity without affecting normal cellular behaviour in monolayers. (n=3) A-B) Internalisation of Cy3-labelled nanoparticles (in red) did not affect proliferative or migratory processes in dermal fibroblasts up to 96 hours post-transfection. C-D) Quantification of metabolic activity and DNA content reveals no cytotoxic outcome from miRNA-29b treatment with varying CR. E-G) Quantification α-SMA expression exhibited a clear intensity decrease with the CR8 (20 pmol) formulation after 7 days without a reduction in cell nuclei.

Having validated the safe intracellular delivery of miRNA nanoparticles, the functional effects of miRNA-29b delivery were assessed by optimising the charge ratio (CR) and miRNA dose. A fibrotic phenotype was first established by treating dermal fibroblasts with TGF-β1, with 10 ng mL^-1^ identified as the optimal concentration to induce an α- SMA-positive myofibroblast phenotype without adversely affecting cell viability (Supp. Fig. 2). Treatment with miRNA-29b nanoparticles formulated at different CRs did not negatively affect cell viability. Instead, a trend towards increasing metabolic activity was observed in miRNA-29b-treated cells relative to untreated controls (Fig. 2C). Consistent with these findings, DNA quantification revealed no significant differences among groups over 7 days post-transfection (Fig. 2D). Quantification of nuclei (Fig. 2E) revealed a higher cell number on day 7 with the CR8 formulation compared to non-treated cells. Importantly, α-SMA quantification in TGF-β1-stimulated fibroblasts showed a ∼2-fold reduction at day 7 following transfection with the CR8 formulation compared to all other groups (Fig. 2F-G). In addition, assessment of miRNA-29b dose demonstrated that the higher dose (40 pmol) produced greater suppression of α-SMA expression without affecting cellular processes (Supp. Fig. 3). Collectively, these findings identify the CR8 formulation containing 40 pmol miRNA-29b as the optimal formulation, demonstrating potent anti-fibrotic activity through the inhibition of myofibroblast differentiation while maintaining normal cell function.

### 3.3. Delivery of optimised miRNA-29b nanoparticles reduces fibrotic markers and modulates matrix deposition in TGF-β1-stimulated fibroblasts

Having determined the optimal miRNA-29b nanoparticle formulation (CR8, 40 pmol) that obtained the highest reduction in α-SMA expression, functional performance was assessed against scrambled mimic nanoparticles (miRNA-Scr). Initially, dermal fibroblast viability was assessed following transfection with miRNA-29b and miRNA-Scr nanoparticles (Fig. 3A). Delivery of miRNA-Scr nanoparticles to TGF-β1- stimulated fibroblasts exhibited a ∼40% decrease in metabolic activity on day 3 post-transfection compared to both non-treated and miRNA-29b-treated cells. Importantly, this response was recovered by day 7 post-transfection. However, DNA quantification did not reveal any apparent differences among groups over 7 days of culture (Fig. 3B).

**Figure 3.**
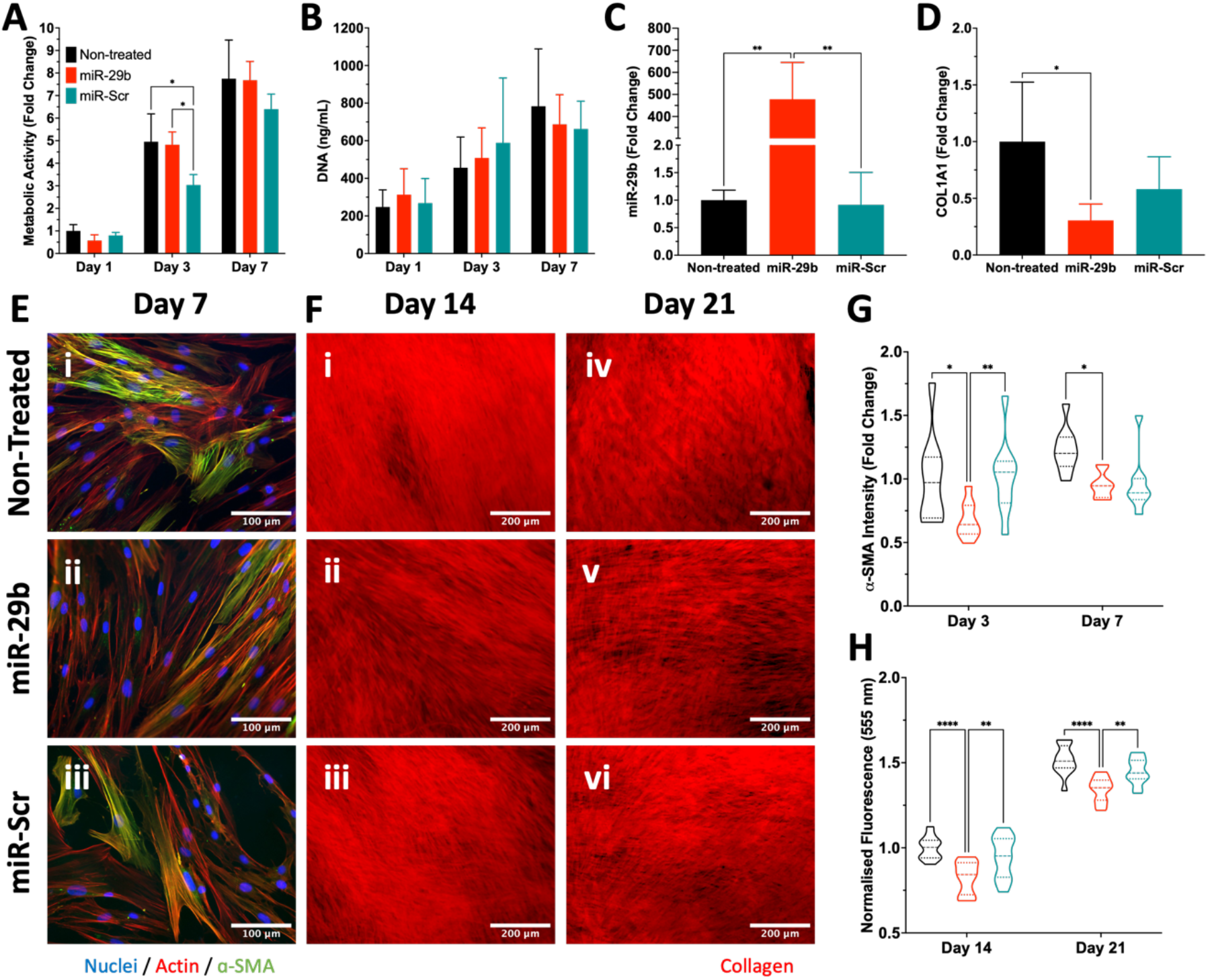
Delivery of miRNA-29b nanoparticles to TGF-β1-stimulated fibroblasts inhibits pro-fibrotic outcomes compared to miRNA-Scr delivery. (n=3) A-B) Assessment of cell viability through metabolic activity and DNA content validates the safety of the miRNA-29b nanoparticle treatment. C-D) Quantification of miRNA-29b expression is highly upregulated following miRNA-29b delivery, leading to a clear downregulation of collagens type I. E-F) Visualisation of α-SMA expression and collagen deposition (555 nm) from TGF-β1-treated fibroblasts exhibit a reduced fibrotic outcome following miRNA-29b treatment. G-H) Quantification of α-SMA intensity and collagen deposition through picrosirius red fluorescence supports previous findings with a clear downregulation of pro-fibrotic markers in the miRNA-29b-treated cells.

Importantly, miRNA-29b nanoparticle delivery enhanced the expression of the target miRNA (Fig. 3C) which led to the downregulation of extracellular matrix components collagen type I (Fig. 3D) and collagen type III (Supp. Fig. 4) by approximately 75% and 30% compared to non-treated cells, respectively. Moreover, quantification of fibronectin 1 expression showed no apparent change among groups. Expression of matrix metalloproteinase-3 (MMP3) showed an increasing trend in nanoparticle-treated fibroblasts while the expression of the TGF-β1 pathway-associated protein SMAD3 was unaffected (Supp. Fig. 4).

Visualisation of TGF-β1-stimulated fibroblasts demonstrated reduced α-SMA expression following miRNA-29b nanoparticle treatment at day 7 (Fig. 3E). These findings were further corroborated by analysis of α-SMA expression, which revealed a ∼20% reduction of α-SMA fluorescence at both 3 and 7 days post-transfection compared with untreated controls (Fig. 3G). Notably, this suppression of α-SMA expression occurred without significant alterations in SMAD2/3 phosphorylation or the expression of the focal adhesion-associated protein vinculin (Supp. Fig. 5), indicating that miRNA-29b selectively attenuated myofibroblast differentiation without disrupting canonical TGF-β1 signalling or cell adhesion.

The anti-fibrotic effects of miRNA-29b delivery were further supported by assessment of collagen deposition using picrosirius red staining, which showed visibly reduced collagen accumulation in treated cells (Fig. 3F). Quantification of picrosirius red fluorescence confirmed a sustained reduction in collagen deposition, with fluorescence intensity decreasing by ∼15% after 14 and 21 days following miRNA-29b treatment (Fig. 3H). Together, these findings demonstrate that miRNA-29b delivery supresses key hallmarks of fibrosis by reducing myofibroblast differentiation and ECM deposition while preserving normal cellular signalling and adhesion.

### 3.4. Nanoparticle incorporation in a collagen-GAG scaffold enables sustained and localised release of miRNA-29b without compromising modulation of target genes

Having established the optimal miRNA-29b formulation capable of inhibiting fibrotic outcomes without compromising normal cellular behaviour, the anti-fibrotic capabilities of the optimised nanoparticles were assessed following incorporation within the collagen-GAG (CG) scaffolds. Initial visualisation of the scaffold microarchitecture through scanning electron microscopy showed that the porous structure was unaffected following miRNA-29b nanoparticle loading (Fig. 4A). Moreover, nanoparticles were distributed homogeneously on the surface while maintaining a rounded morphology (Fig. 4B-C). Subsequent characterisation of nanoparticle release kinetics showed a gradual release of up to 4% of the initial cargo into the surrounding medium, suggesting that most of the nanoparticles remain within the scaffold facilitating localised cellular uptake (Fig. 4D).

**Figure 4.**
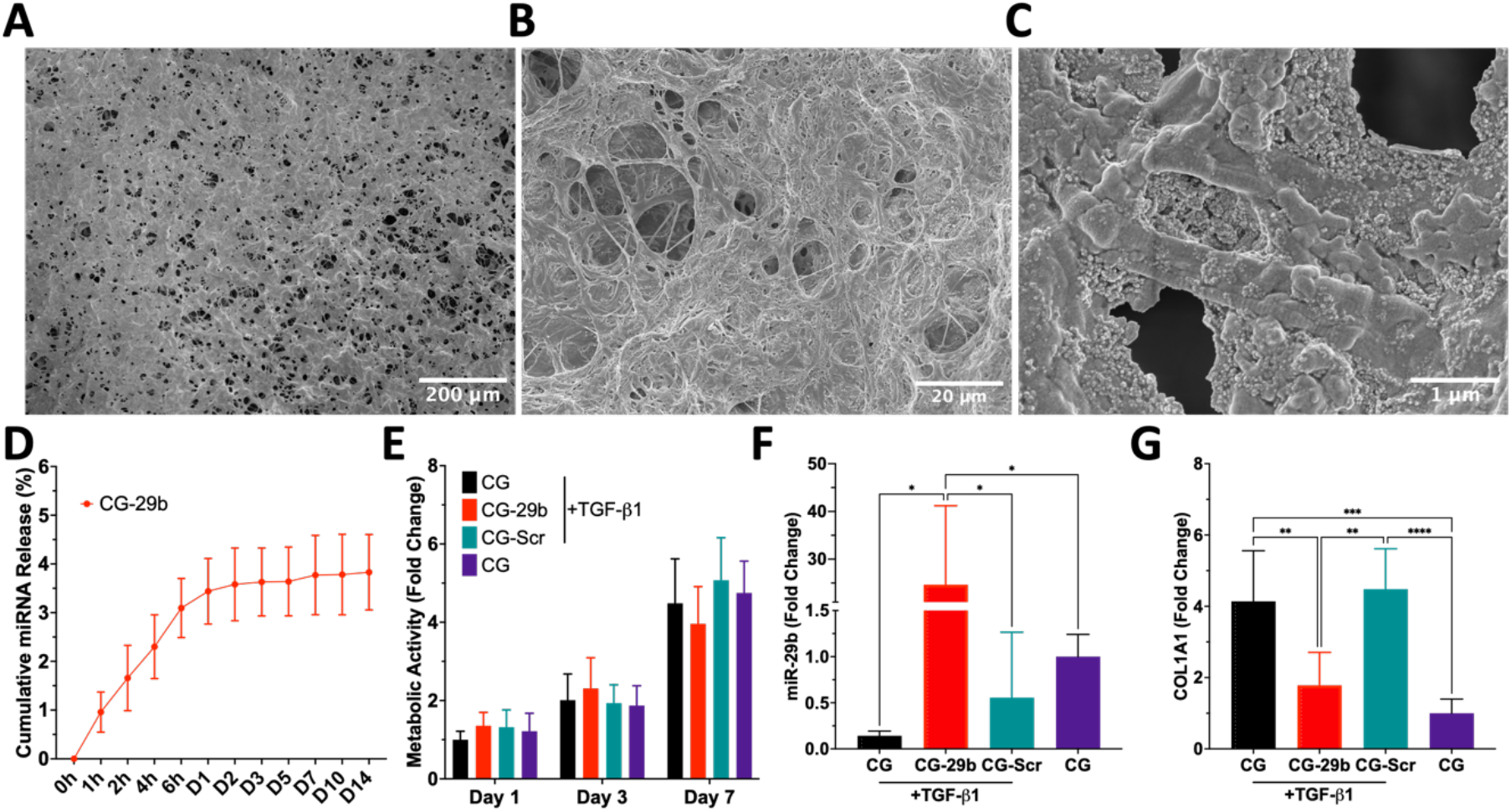
CG-29b scaffolds promote downregulation of fibrotic genes following gradual localised release of miRNA-29b. (n=3) A-C) SEM visualisation of microarchitecture shows no changes in porosity while nanoparticles maintain a rounded morphology. D) Analysis of release kinetics revealed the gradual release of miRNA-29b nanoparticles up to 14 days. E) Characterisation of cell viability through metabolic activity showed no apparent differences among groups. F) Expression of target miRNA-29b was effectively upregulated following miRNA-29b nanoparticle transfection. G-I) Scaffold-mediated miRNA-29b delivery decreased the excessive expression of collagen type I compared to TGF-β1-treated cells on CG only and CG- Scr scaffolds.

Following nanoparticle characterisation, the biological and functional outcomes of scaffold-mediated miRNA-29b delivery were evaluated. Cell viability, assessed by metabolic activity, showed no apparent differences between the CG-29b group and the controls, regardless of TGF-β1-supplementation, up to 7 days post-transfection (Fig. 4E). Quantification of miRNA-29b expression exhibited a ∼20-fold upregulation of the target miRNA on the CG-29b scaffold groups compared to all other TGF-β1 supplemented and not-supplemented groups (Fig. 4F). Importantly, scaffold-mediated miRNA-29b upregulation resulted in a reduction of collagen type I expression relative to TGF-β1-treated cells on scaffolds (Fig. 4G), while expression of collagen type III and fibronectin-1 remained unchanged (Supp. Fig. 6). This selective modulation is notable, as fibronectin plays a critical role in provisional matrix formation and stabilisation, processes that influence wound healing outcomes and the progression of fibrosis.^52^ Furthermore, MMP3 expression was restored in the CG-29b scaffold groups to levels comparable to those observed in the non-fibrotic control (Supp. Fig. 6), suggesting that miRNA-29b delivery promotes a shift towards physiological ECM remodelling.

### 3.5. Scaffold-mediated miRNA-29b delivery mitigates matrix contraction and **α-SMA expression**

Having established that scaffold-mediated miRNA-29b delivery effectively modulated anti-fibrotic gene expression in TGF-β1-stimulated dermal fibroblasts, a collagen gel contraction assay was performed to evaluate miRNA-29b’s effects on myofibroblast-mediated matrix contraction. Visual inspection of collagen gels containing miRNA-29b nanoparticles showed reduced contraction compared with both miRNA-Scr and non-treated controls under fibrotic conditions (Fig. 5A). Notably, the extent of gel contraction in the miRNA-29b group closely resembled that of non-treated gels without TGF-β1 supplementation, suggesting that miRNA-29b attenuated the differentiation into the contractile myofibroblast phenotype. Quantitative analysis confirmed these observations, with the miRNA-29b group exhibiting contraction levels comparable to the non-fibrotic control throughout the 7-day culture period despite exposure to the pro-fibrotic stimulus (Fig. 5B).

**Figure 5.**
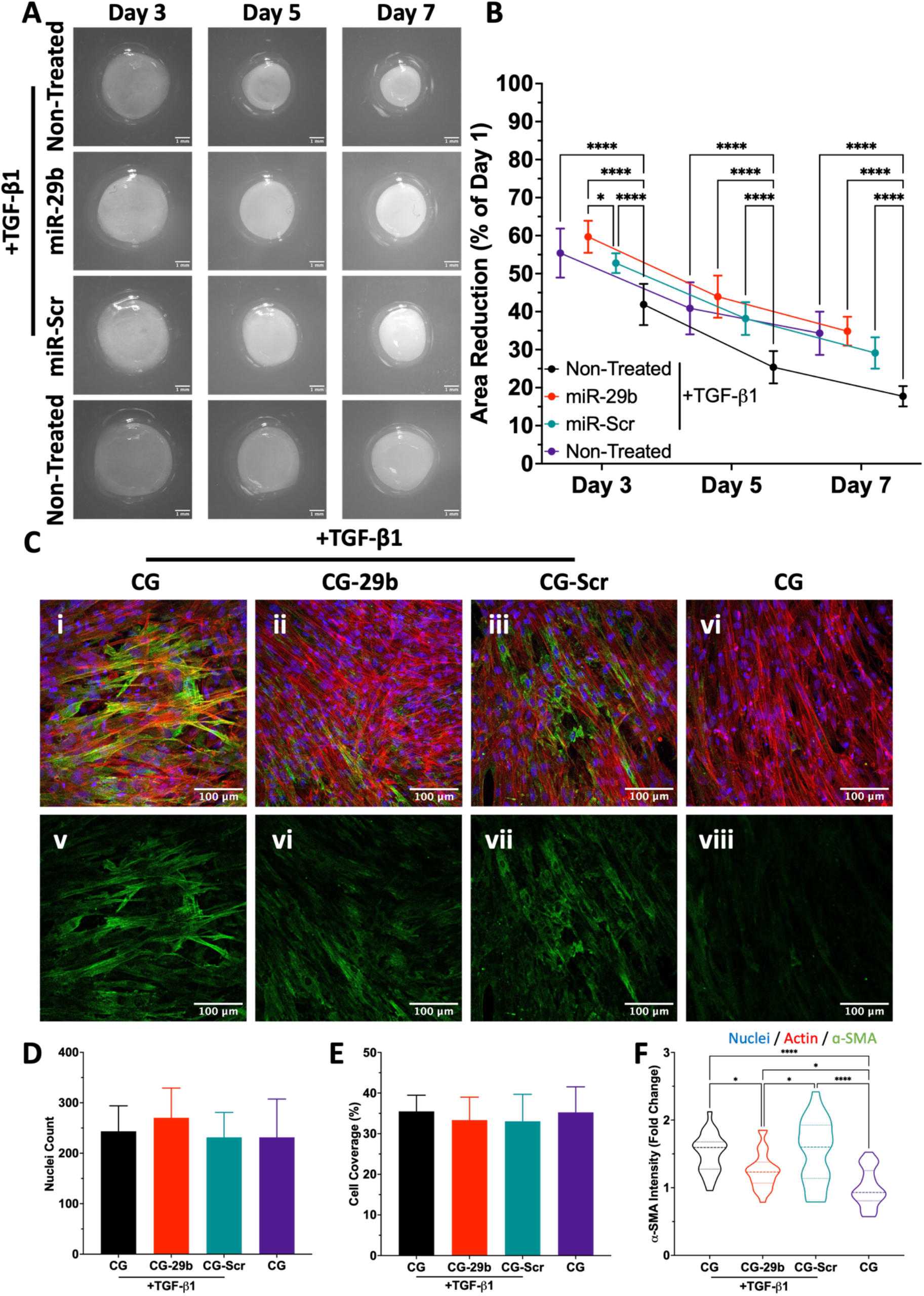
CG-29b scaffolds mitigate fibrotic responses following gradual localised release of miRNA-29b. (n=4) A) Analysis of matrix contraction showed similar area reduction in the miRNA-29b group following TGF-β1 supplementation compared to the non-fibrotic control. B) Quantification of relative area reduction in each group exhibits an enhanced anti-fibrotic outcome following miRNA-29b delivery comparable to non-fibrotic conditions. C) Visualisation and quantification of nuclei count, cell coverage and α-SMA intensity reveal a clear inhibition of myofibroblast differentiation in CG-29b scaffolds, trending towards a non-fibrotic state.

To further assess the anti-fibrotic effects of scaffold-mediated miRNA-29b delivery, cell morphology and α-SMA expression were evaluated in TGF-β1-conditioned dermal fibroblasts cultured on miRNA-activated scaffolds. Cells cultured on CG-29b scaffolds retained a predominantly spindle-shaped morphology and exhibited reduced α-SMA expression compared with fibrotic control groups (Fig. 5C). Quantification of nuclei number (Fig. 5D) and cell coverage (Fig. 5E) revealed no apparent differences between groups, indicating that the observed reductions in expression were not attributable to changes in cell number or scaffold colonisation. Consistent with these findings, α-SMA fluorescence intensity in the CG-29b group remained only ∼20% higher than that of the non-fibrotic CG control (Fig. 5F). In contrast, α-SMA expression in the TGF-β1-treated CG and CG-Scr scaffold groups was ∼25% greater than in the CG-29b group, demonstrating that scaffold-mediated miRNA-29b delivery substantially attenuated TGF-β1-induced myofibroblast differentiation while preserving cell viability.

### 3.6. Scaffold-mediated miRNA-29b delivery prevents excessive ECM deposition in pro-fibrotic conditions

After validating that scaffold-mediated miRNA-29b delivery to TGF-β1-stimulated myofibroblasts reduced α-SMA expression and matrix contraction, we next investigated whether miRNA-29b upregulation also influence ECM deposition dynamics within the 3D scaffold microenvironment. However, assessing cell-associated ECM deposition proved challenging, as conventional histological and fluorescence-based techniques were unable to distinguish newly synthesised ECM from the CG scaffold template. To overcome this limitation, we adapted a metabolic labelling approach previously described by Loebel et al.^49^, in which nascent cell-deposited ECM is labelled using the azide-containing methionine analogue L-azidohomoalanine (AHA). Following fluorescent conjugation, this strategy enabled selective visualisation of newly deposited ECM while distinguishing it from the underlying CG scaffold.

Visualisation of ECM deposition over 14 days revealed marked differences between the CG-29b scaffolds and the other TGF-β1-treated groups (CG and CG-Scr). Despite continuous exposure to the profibrotic stimuli, the CG-29b scaffolds exhibited reduced cellular coverage and ECM deposition (Fig. 6A). Furthermore, compared with non-fibrotic CG scaffolds cultured without TGF-β1 supplementation, only the CG-29b groups demonstrated comparable cell density and ECM deposition, particularly at day 7, although this similarity became less pronounce by day 14.

**Figure 6.**
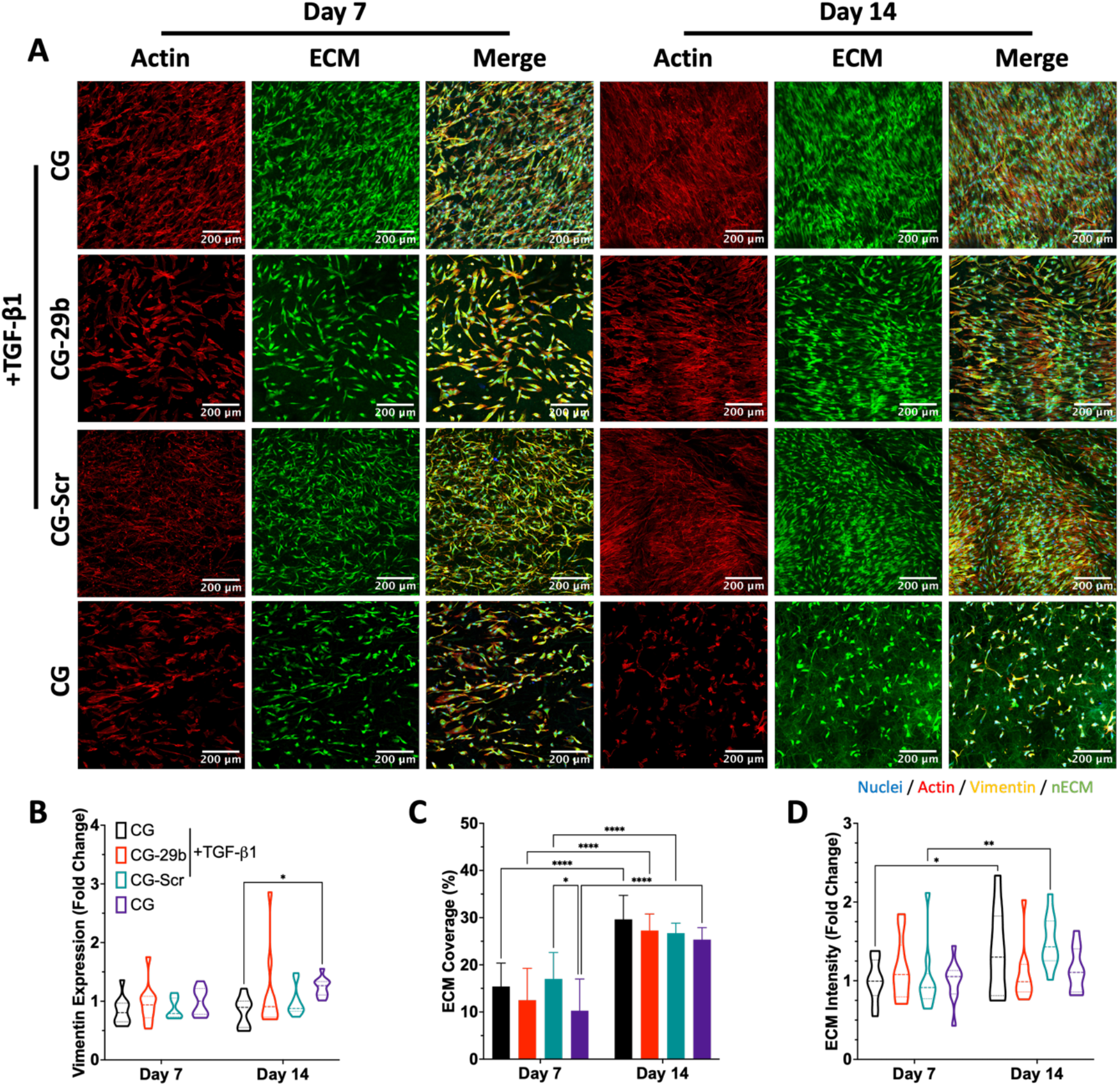
CG-29b scaffolds promote a matrix deposition profile comparable to non-fibrotic conditions despite the presence of pro-fibrotic stimuli. (n=5) A) Fluorescent visualisation of ECM and cell coverage revealed decrease cell number and ECM expression in the CG-29b and non-fibrotic control. B) Quantification of vimentin expression displayed increased intensity in the non-fibrotic groups relative to the TGF-β1-treated CG scaffolds. C-D) Analysis of ECM coverage and intensity showed a comparable behaviour between the CG-29b group and the non-fibrotic control with reduced ECM coverage and intensity on days 7 and 14, respectively.

Semi-quantitative analysis of vimentin expression showed significantly greater protein expression in the non-fibrotic control than in the TGF-β1-stimulated CG group at day 14 (Fig. 6B, Supp. Fig. 7), whereas no appreciable differences were observed after 7 days of culture. A similar trend was observed for ECM coverage, with the CG-29b scaffolds closely resembling the non-fibrotic control at day 7 despite its increase for all groups at day 14 (Fig. 6C). Consistent with these findings, quantification of ECM fluorescence intensity showed a trend towards increased signal in the TGF-β1 treated CG-Scr group relative to both the CG-29b scaffolds (p < 0.06) and the non-fibrotic control (p < 0.13) (Fig. 6D). Together, these results suggest that miRNA-29b delivery attenuated ECM deposition in the 3D microenvironment, maintaining a matrix deposition profile more comparable to the non-fibrotic condition despite the presence of TGF-β1 and validating the results obtained in 2D monolayer cultures.

## 4. Discussion

The primary objective of this study was to develop a miRNA-29b mimic-activated scaffold capable of mitigating fibrotic outcomes during wound healing by reducing of α-SMA-mediated matrix contraction and excessive ECM deposition. Initially, the role of miRNA-29b in ECM-associated processes was validated through bioinformatic analysis, followed by successful cellular internalisation of GET-based nanoparticles without detrimental effects on fibroblast viability. The anti-fibrotic efficacy of the optimised miRNA-29b nanoparticles was subsequently demonstrated by significant reductions in collagen deposition and α-SMA expression from TGF-β1-stimulated fibroblasts. The optimised miRNA-29b formulation was then incorporated into collagen-GAG (CG) scaffolds, which effectively modulated fibrotic gene expression while preserving a scaffold structure conducive to fibroblast infiltration and proliferation. Finally, functional outcomes, including reduced matrix contraction, α- SMA expression, and ECM deposition, were comparable to those observed in non-fibrotic conditions, thereby confirming the therapeutic potential of scaffold-mediated miRNA-29b delivery. Taken together, these findings demonstrate that scaffold-mediated miRNA-29b delivery represents a promising anti-fibrotic strategy for wound healing by reducing matrix contraction, α-SMA expression, and excessive ECM deposition.

Fibrotic responses during wound healing are primarily driven by the TGF-β signalling pathway. Under physiological conditions, TGF-β promotes the differentiation of fibroblasts into myofibroblasts, generating highly active and contractile cells through the canonical SMAD2/3 signalling pathway, which leads to upregulation of α-SMA.^53,54^ In vitro exposure of dermal fibroblasts to TGF-β1 resulted in a dose-dependent upregulation of α-SMA, confirming the establishment of a myofibroblast phenotype consistent with published literature and providing a suitable *in vitro* model for investigating the anti-fibrotic effects of miRNA-29b.^55,56^

MiRNA-29b is a well-documented suppressor of SMAD2/3-mediated fibrosis and is particularly associated with the downregulation of collagen type I deposition.^37^ This role was further supported by predictive bioinformatic analyses using publicly available datasets and software. Additionally, the physicochemical properties of the miRNA-29b nanoparticles were optimised to enhance cellular uptake. MiRNA-29b-loaded GET nanoparticles were engineered with diameters below 200 nm and surface charges exceeding +10 mV, characteristics considered favourable for suitable cellular internalisation.^57,58^ These parameters are known to facilitate uptake via clathrin-mediated endocytosis.^59–61^ Fluorescent labelling confirmed perinuclear localisation of the nanoparticles, indicating successful cellular internalisation. Importantly, no adverse effects on cell proliferation or migration were observed, consistent with previous findings on the biocompatibility of GET-based delivery systems.^21,28,43,62^ Among the formulations tested, the CR8 formulation (123.2 nm, > +20 mV) administered at a dose of 40 pmol miRNA produced the most pronounced anti-fibrotic effect, significantly reducing α-SMA expression in TGF-β1-stimulated fibroblasts.

Physiological wound healing requires ECM deposition; however, the quality and composition of the deposited matrix critically influence the risk of pathological scarring and impaired healing.^63–65^ In fibrotic conditions, the balance between ECM deposition and remodelling shifts towards excessive accumulation of fibrillar collagens, particularly types I and III, fibronectin overexpression, and impaired ECM turnover resulting from an imbalance between MMPs and their inhibitors (TIMPs).^64,66^ Treatment with the optimised miRNA-29b nanoparticles reduced the expression of fibrillar collagens and increased MMP-3 expression at the post-transcriptional level, without altering fibronectin or SMAD3 expression relative to controls. Notably, this expression profile confirms that miRNA-29b treatment does not interfere with normal wound healing processes associated with fibronectin-mediated matrix deposition and stabilisation, which are essential for formation of the provisional matrix that determines wound fate.^52^ Instead, miRNA-29b appears to selectively limit the excessive collagen accumulation that drives pathological scar maturation.

Beyond ECM production, α-SMA and focal adhesion proteins such as vinculin contribute the mechanical stabilisation of cell-ECM interactions.^67,68^ *In vitro*, miRNA- 29b delivery effectively normalised α-SMA levels to those observed in non-fibrotic controls, without affecting SMAD2/3 phosphorylation or vinculin expression. These findings indicate that miRNA-29b selectively attenuated myofibroblast differentiation without disrupting canonical TGF-β1 signalling or cell adhesion. Additionally, miRNA-Scr-loaded nanoparticles produced similar effects in some of the analysed genes and proteins; however, these effects were not consistently observed across all analyses, suggesting that they are more likely attributable to the scrambled miRNA sequence itself rather than off-target activity of the GET peptide.^69–71^ Collectively, these findings support the anti-fibrotic potential of the miRNA-29b platform by targeting core drivers of fibrosis while preserving ECM remodelling processes essential for effective wound healing. This selective activity suggests a therapeutic window in which core fibrotic mechanisms are suppressed, yet essential ECM remodelling is preserved.

Collagen-based scaffolds are widely used to treat wounds of diverse aetiologies, with several collagen-based wound dressings already available clinically.^24–27^ These biomaterials provide a structural template that supports cellular infiltration and tissue remodelling which has been shown to limit myofibroblast activity and contraction, thereby enhancing wound healing outcomes.^30^ Furthermore, their ease of functionalisation enables the incorporation of advanced therapeutic strategies, including gene delivery, to further promote tissue regeneration.^28^ Following optimisation of the anti-fibrotic miRNA-29b formulation, the nanoparticles were successfully incorporated into the CG scaffold. Importantly, this modification did not alter the scaffold’s porous microarchitecture, preserving the microenvironment required for cell infiltration, proliferation, and tissue regeneration.^72^ The incorporated nanoparticles retained uniform surface distribution and spherical morphology, consistent with previous findings.^21^ Moreover, the functionalised scaffold enabled sustained, localised delivery of miRNA-29b over 14 days, a release profile well suited to modulating early fibrotic responses while supporting long-term tissue remodelling.

Functionalisation of the CG scaffold with the miRNA-29b delivery system imparted anti-fibrotic activity while maintaining the scaffold’s regenerative properties. The platform supported cell viability and proliferation, and gene expression analysis confirmed successful miRNA-29b upregulation, although expression levels were lower than those observed in monolayer cultures. Despite this reduction, scaffold-mediated miRNA-29b delivery is likely to provide greater therapeutic durability than direct nanoparticle administration due to its sustained release profile. This prolonged and localised delivery may also minimise off-target effects while enhancing long-term anti-fibrotic efficacy. Functionally, scaffold-mediated miRNA-29b delivery effectively downregulated collagen type I expression to levels comparable with non-fibrotic conditions, indicating robust anti-fibrotic activity. Together, these findings demonstrate a shift from pathological ECM accumulation towards physiological remodelling, even in the presence of fibrotic stimuli, highlighting both the versatility of the scaffold as a gene delivery platform and the importance of a regenerative ECM.

Persistent collagen deposition and myofibroblast expansion drive progressive tissue stiffening, scar formation, and functional impairment.^5^ Scaffold-mediated miRNA-29b delivery significantly attenuated myofibroblast-mediated matrix contraction, reduced α-SMA expression, and restored ECM deposition towards a non-fibrotic fibroblast phenotype. Both matrix contraction and ECM accumulation were markedly reduced, suggesting decreased mechanical strain that may help preserve tissue architecture and prevent contracture- and stiffening-related complications. Such pathological changes can ultimately lead to permanent tissue shortening, impaired mobility, pain, and long-term disability, making this a more effective scaffold platform for mitigating fibrosis.^73–75^ In contrast, fibrotic fibroblasts cultured on miRNA-free scaffolds maintained elevated α-SMA expression and continued to deposit excessive ECM. These findings are consistent with prior studies demonstrating that collagen scaffold-based delivery of miRNA-29b suppresses collagen synthesis and limits wound contraction.^76^ The present study further demonstrates enhanced anti-fibrotic efficacy across genetic, molecular, and functional levels, supporting the potential of the scaffold as a simple yet effective approach to anti-fibrotic wound healing, with advantages over comparable strategies reported in the literature.^77,78^

Additionally, the translational potential of scaffold-mediated miRNA-29b delivery is strengthened by limitations associated with current gene therapy strategies. Although numerous gene delivery systems exhibit promising anti-fibrotic effects, many are limited by burst release, inadequate localisation, and subsequent off-target activity.^79^ Consequently, increasing attention has focused on targeting multiple miRNAs to regulate fibrotic pathways, yet most systems lack matrix-mediated control over their release. For instance, inhibiting miRNA-21 can reduce fibrotic markers and tissue contraction, thereby attenuating scar formation.^80,81^ However, miRNA-21 also supports angiogenesis, cell proliferation, immuno-modulation, and re-epithelialisation; thus, its inhibition may inadvertently delay wound healing.^82,83^ Similarly, inhibition of miRNA- 182 reduces collagen type I and fibronectin expression via TGF-β1/SMAD modulation but could aggravate inflammation due to its known anti-inflammatory roles in ischemic injuries.^84–87^ Supplementation of miRNA-9 can counteract fibrosis and inflammation, yet its involvement in neurogenesis raises concerns about disrupting neural repair in skin and surrounding tissues given the synergistic role of neurons and endothelial cells crosstalk in wound healing.^88,89^ These reports underscore the challenges of systemic or poorly controlled miRNA delivery, further emphasising the advantages of our scaffold-mediated miRNA-29b delivery system as a platform for sustained, localised, and targeted therapy.

Increasingly, there is consensus that combining biomaterial scaffolds with gene delivery offers the most promising strategies for treating wound healing-associated pathologies including fibrotic conditions, due to their complex microenvironment.^3,79^ In this context, scaffold-mediated miRNA-29b delivery represents a clinically relevant and practical solution, offering localised and sustained anti-fibrotic therapy while addressing key limitations of conventional therapeutic approaches. Beyond cutaneous healing, scaffold-mediated miRNA-29b delivery may also be applicable to multiple conditions involving pulmonary, hepatic, and renal systems due to the wide influence of miRNA-29b in fibrosis. In summary, scaffold-mediated miRNA-29b delivery offers a modern, targeted, and potentially safer alternative for mitigating fibrosis, minimising scar formation, and improving overall wound healing outcomes.

## 5. Conclusion

Overall, this work outlines the development of collagen-based scaffolds functionalised with a miRNA-29b delivery system to promote anti-fibrotic responses via reduced α- SMA expression, matrix contraction, and excessive ECM accumulation for wound healing applications. Bioinformatic analysis validated the role of miRNA-29b in ECM-associated processes and miRNA-29b nanoparticles displayed suitable properties to mitigate fibrotic outcomes in TGF-β1-stimulated dermal fibroblasts. Delivery of miRNA-29b nanoparticles resulted in reduced collagen deposition and α-SMA expression. Moreover, development of miRNA-29b-activated scaffolds elicited a similar genetic profile to exogenous delivery of the miRNA while allowing for the sustained and localised release of the cargo. Finally, miRNA-29b-activated scaffolds reduced matrix contraction and ECM deposition to levels comparable with non-fibrotic conditions. Taken together, these findings demonstrate the potential of the miRNA- 29b-activated scaffold as a promising anti-fibrotic platform which provides a toolbox of options for treating wound healing-associated pathologies.

## Author Contributions

JCPC (data curation, validation, formal analysis, visualisation, investigation, methodology, and writing – original draft), AE (investigation, validation, and data curation), MD (investigation, validation, methodology, and writing – review and editing), AA (investigation, validation, and data curation), JM (investigation and writing – review and editing), JED (resources), CJK (conceptualization, supervision, and writing – review and editing), SB (conceptualization, supervision, and writing – review and editing), FOB (conceptualization, supervision, funding acquisition, and writing – review and editing).

## Conflicts of Interest

No conflicts to declare

## Data Availability

Data for this article are openly available at Open Science Framework (OSF) Repository at under the terms of the Creative Commons Attribution 4.0 (CC-BY 4.0) license.

## Acknowledgements

The authors acknowledge the Research Ireland Advanced Materials and Bioengineering Research (AMBER) Centre for providing financial support (SFI/12/RC/2278_P2). The authors would also like to acknowledge financial support from Debra Ireland (22563A01), the Higher Education Authority (HEA), the Department of Further and Higher Education, Research, Innovation and Science (DFHERIS), the Shared Island Fund through the North South Research Program, the Health Research Board (HRB) Investigator-Led Projects (ILP) 2024 (ILP-POR-2024- 064), and HORIZON EUROPE under the WIDERA Twinning Programme (Grant 101079123; ‘REGENEU’). JED would like to acknowledge funding by the Defence Accelerator (DASA, DSTL) (Award reference: ACC6007330), the European Research Council under the European Community’s Seventh Framework Programme (FP7/2007–2013)/ERC grant agreement 227845, the Medical Research Council (grant number MR/K026682/1); the Engineering and Physical Sciences Research Council; and the Biotechnology and Biological Sciences Research Council, for the UK Regenerative Medicine Platform Hub “Acellular Approaches for Therapeutic Delivery”. Additionally, the authors would like to acknowledge the support given by Dr Brenton Cavanagh. Collagen materials were provided by Integra Life Sciences, Inc. through a Material Transfer Agreement.

## Notes

### Competing Interest Statement

The authors have declared no competing interest.

