## Supplementary Files for "Scaffold-mediated delivery of a miRNA-29b mimic mitigates excessive extracellular matrix deposition and matrix contraction in wound healing applications"

### Supplementary Tables

**Supplementary Table 1.** Lists of genes analysed by qRT-PCR.

| Target Protein | Target Gene Reference | GeneGlobeID |
|---|---|---|
| **18S Ribosomal RNA (18S)** | Hs_RRN18S_1_SG | QT00199367 |
| **Collagen Type I** | Hs_COL1A1_1_SG | QT00037793 |
| **Collagen Type III** | Hs_COL3A1_1_SG | QT00058233 |
| **Matrix Metalloproteinase 3 (MMP-3)** | Hs_MMP3_1_SG | QT00060025 |
| **Small Mother Against Decapentaplegic (SMAD) protein 3** | Hs_SMAD3_1_SG | QT00008729 |
| **Fibronectin 1** | Hs_FN1_1_SG | QT00038024 |
| **microRNA-29b** | hsa-miR-29b-3p | 478369_mir |

### Supplementary Figures

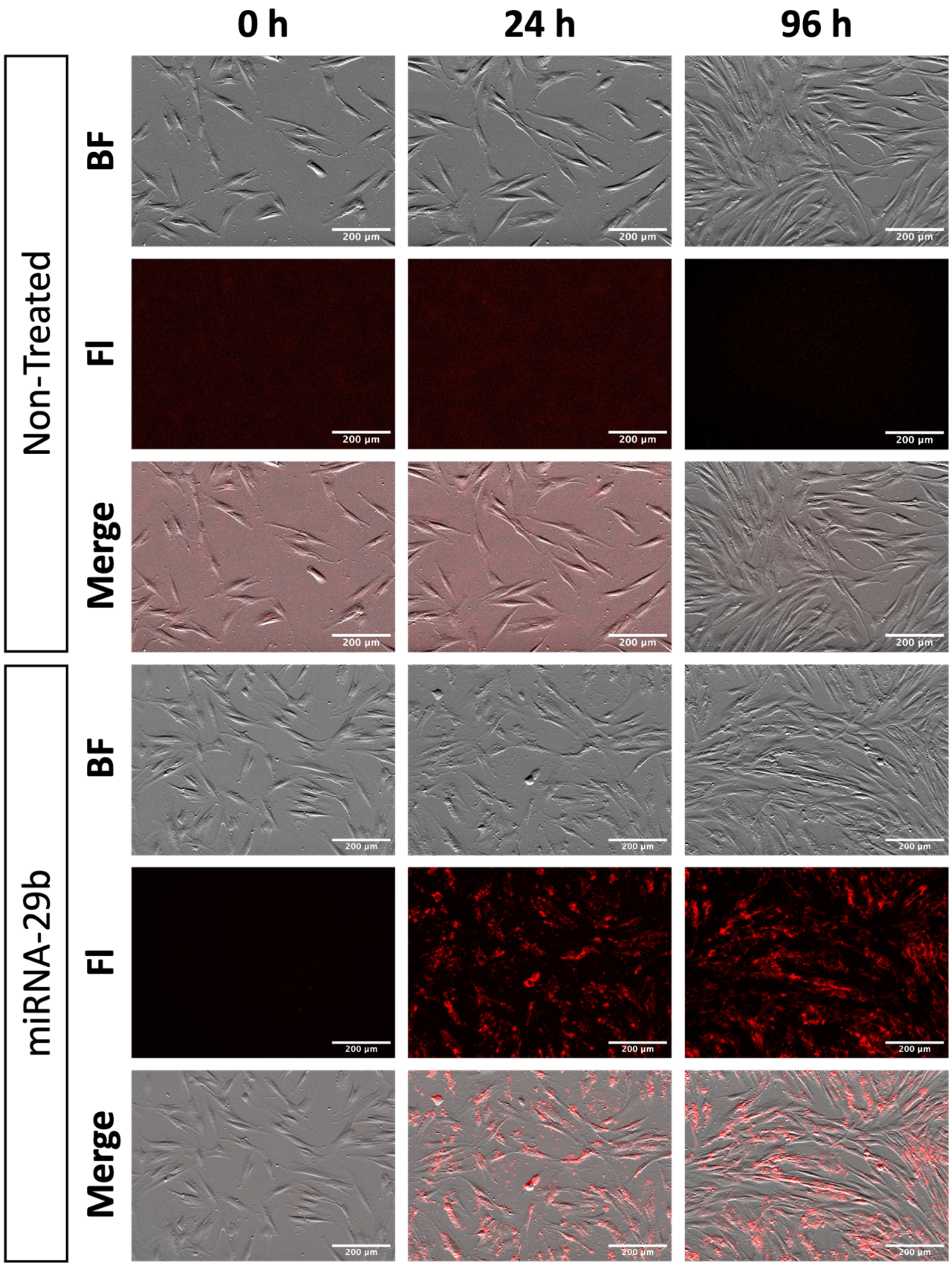


**Supplementary Figure 1. miRNA nanoparticles are effectively internalised in dermal fibroblasts without a detrimental effect on cell migration or proliferation.**

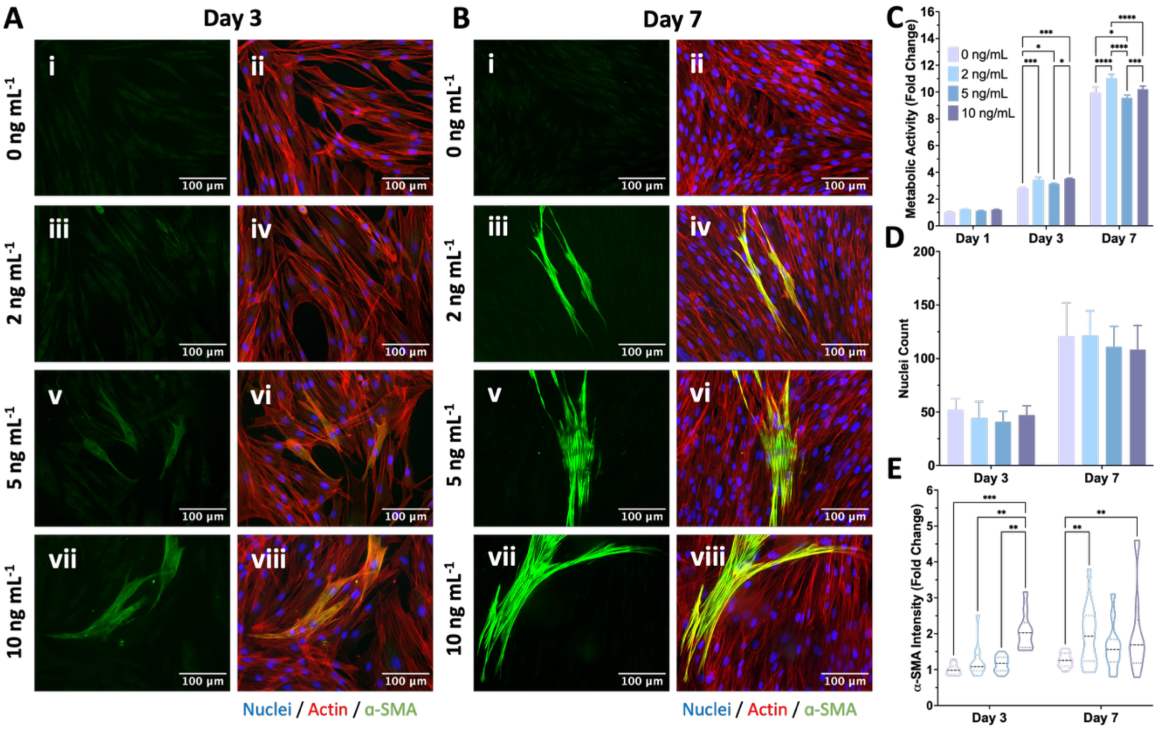


**Supplementary Figure 2. Expression of α–SMA is significantly increased in dermal fibroblasts on days 3 and 7 post-transfection when conditioning cells with TGF-β1.** (n=3) A-B) Representative images of TGF-β1-conditioned dermal fibroblasts show a dose-dependent differentiation over 7-days C). Assessment of metabolic activity does not display a detrimental effect on the fibroblasts when treated with varying TGF-β1 concentrations. D) Quantification of cell nuclei shows no differences among different doses. E). Analysis of α-SMA expression from TGF-β1-treated fibroblasts reveals that 10 ng $mL^{-1}$ significantly increases the expression of the marker after 3 and 7 days. Data shows mean ± SD, * indicates $p<0.05$, ** $p<0.01$, *** $p<0.001$, and **** $p<0.0001$.

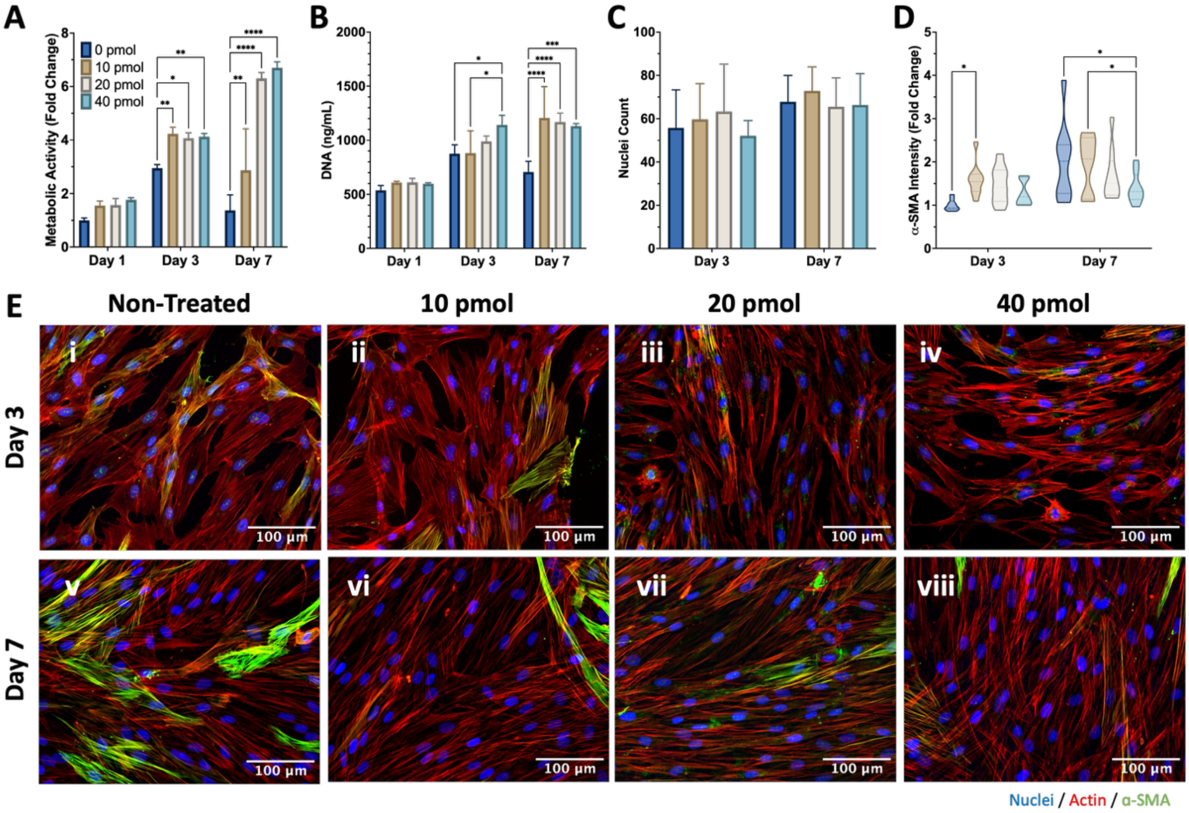


**Supplementary Figure 3. Delivery of miRNA-29b nanoparticles with varying dose reduces α-SMA expression in TGF-β1-conditioned dermal fibroblasts**. (n=3) A-B) Quantification of metabolic activity and DNA content reveals no cytotoxic outcome from miRNA-29b treatment with varying dose. C-E) Quantification of nuclei number and α-SMA expression exhibited a significant decrease relative to the negative control with the highest dose after 7 days. Data shows mean ± SD, * indicates $p<0.05$, ** $p<0.01$, *** $p<0.001$, and **** $p<0.0001$.

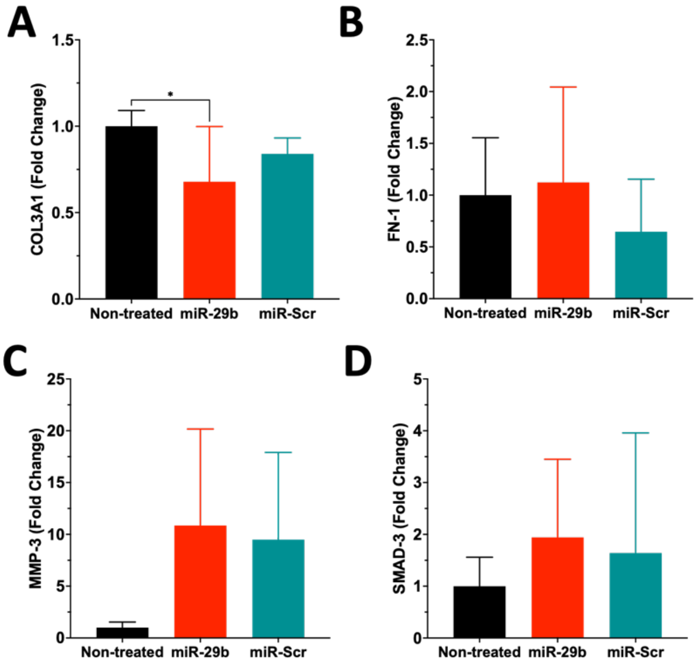


**Supplementary Figure 4. Delivery of miRNA-29b downregulates collagen type III in TGF-β1-conditioned dermal fibroblasts.** (n=6) A-B) Expression of downstream genes of interest exhibit a clear downregulation of collagens type III after miRNA-29b treatment without affecting the expression of other ECM proteins like fibronectin 1. C-D) Gene expression of MMP3 was upregulated in nanoparticle-treated groups while SMAD3 was unchanged. Data shows mean ± SD, * indicates $p<0.05$.

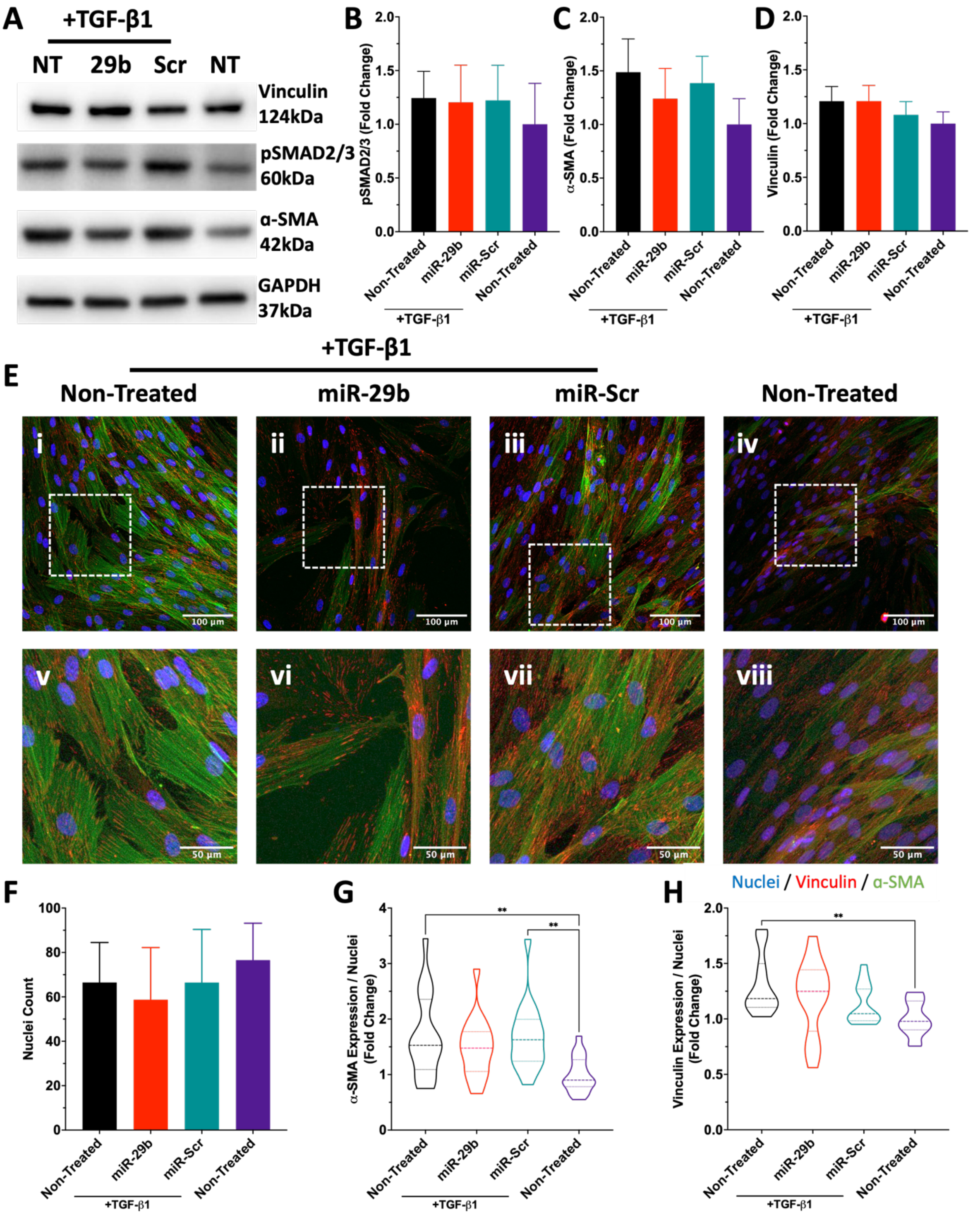


**Supplementary Figure 5. Delivery of miRNA-29b does not affect SMAD2/3 phosphorylation or vinculin expression**. (n=4) A-D) Characterisation and quantification of phosphorylated SMAD2/3, α-SMA, and vinculin protein expression following miRNA-29b delivery. E) Visualisation of α-SMA and vinculin expression shows evident co-localisation of the proteins. F-H) Delivery of miRNA-29b reduces α-SMA expression but does not affect vinculin intensity. Data shows mean ± SD, ** indicates p<0.01.

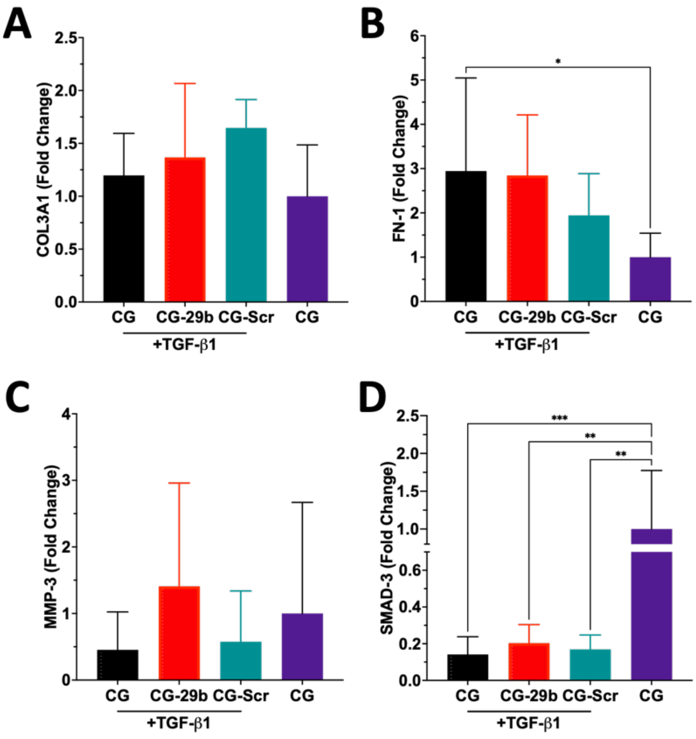


**Supplementary Figure 6. Scaffold-mediated delivery of miRNA-29b does not affect the expression of collagen type III and fibronectin**. (n=6) A-B) Scaffold-mediated miRNA-29b delivery did not significantly affect collagen type III and fibronectin-1 expression. C-D) Expression of MMP3 showed an increasing trend while SMAD3 was downregulated in CG-29b scaffolds compared to the non-fibrotic control. Data shows mean ± SD, * indicates $p<0.05$, ** $p<0.01$, and *** $p<0.001$.

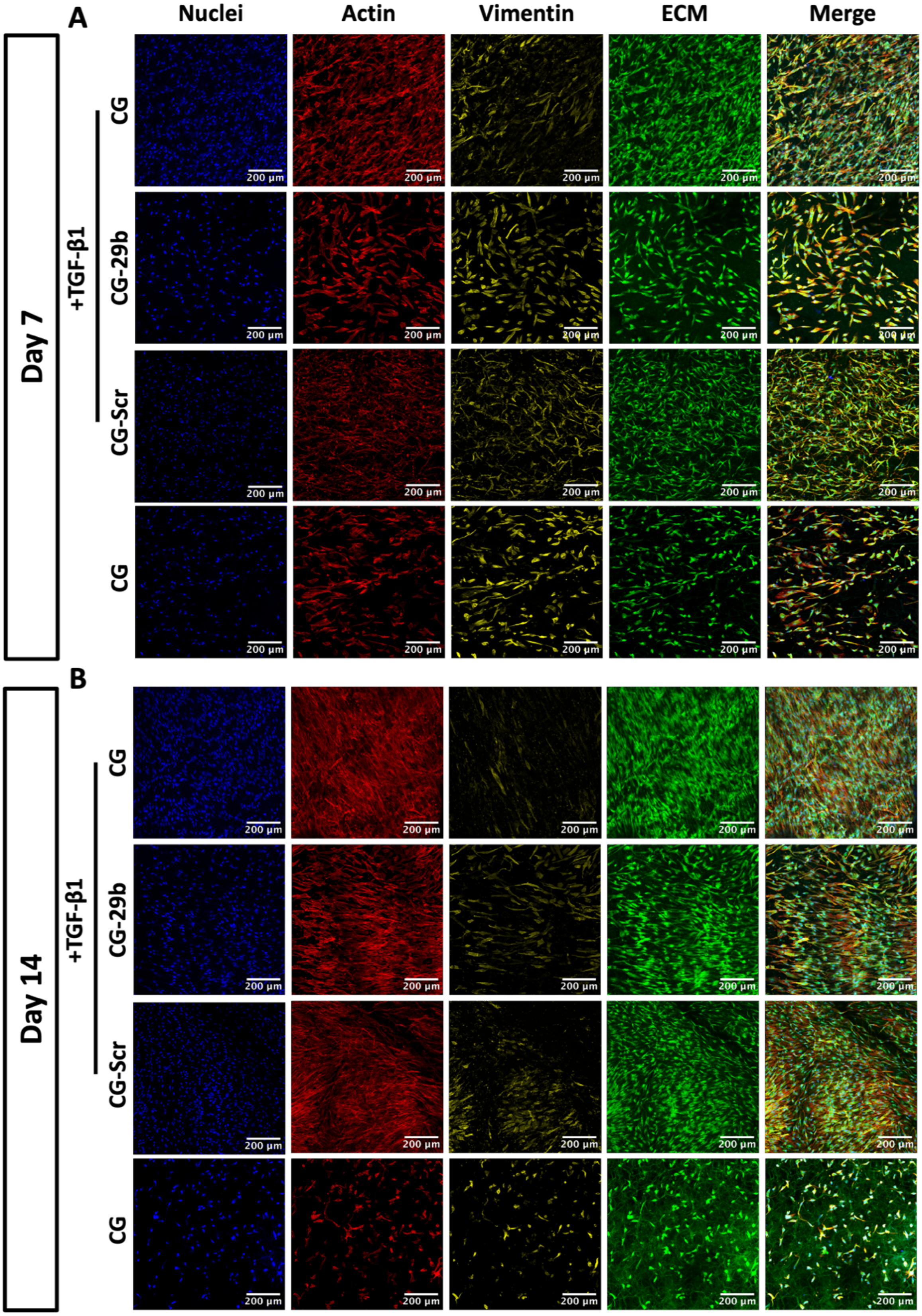


**Supplementary Figure 7. ECM deposition is markedly reduced from TGF-β1-conditioned fibroblasts cultured on CG-29b scaffolds.** (n=4)
